# Kappa Opioid Receptor Antagonism Restores Phosphorylation, Trafficking and Behavior induced by a Disease Associated Dopamine Transporter Variant

**DOI:** 10.1101/2023.05.03.539310

**Authors:** Felix P. Mayer, Adele Stewart, Durairaj Ragu Varman, Amy E. Moritz, James D. Foster, Anthony W. Owens, Lorena B. Areal, Raajaram Gowrishankar, Michelle Velez, Kyria Wickham, Hannah Phelps, Rania Katamish, Maximilian Rabil, Lankupalle D. Jayanthi, Roxanne A. Vaughan, Lynette C. Daws, Randy D. Blakely, Sammanda Ramamoorthy

## Abstract

Aberrant dopamine (DA) signaling is implicated in schizophrenia, bipolar disorder (BPD), autism spectrum disorder (ASD), substance use disorder, and attention-deficit/hyperactivity disorder (ADHD). Treatment of these disorders remains inadequate, as exemplified by the therapeutic use of d-amphetamine and methylphenidate for the treatment of ADHD, agents with high abuse liability. In search for an improved and non-addictive therapeutic approach for the treatment of DA-linked disorders, we utilized a preclinical mouse model expressing the human DA transporter (DAT) coding variant DAT Val559, previously identified in individuals with ADHD, ASD, or BPD. DAT Val559, like several other disease-associated variants of DAT, exhibits anomalous DA efflux (ADE) that can be blocked by d-amphetamine and methylphenidate. Kappa opioid receptors (KORs) are expressed by DA neurons and modulate DA release and clearance, suggesting that targeting KORs might also provide an alternative approach to normalizing DA-signaling disrupted by perturbed DAT function. Here we demonstrate that KOR stimulation leads to enhanced surface trafficking and phosphorylation of Thr53 in wildtype DAT, effects achieved constitutively by the Val559 mutant. Moreover, these effects can be rescued by KOR antagonism of DAT Val559 in *ex vivo* preparations. Importantly, KOR antagonism also corrected *in vivo* DA release as well as sex-dependent behavioral abnormalities observed in DAT Val559 mice. Given their low abuse liability, our studies with a construct valid model of human DA associated disorders reinforce considerations of KOR antagonism as a pharmacological strategy to treat DA associated brain disorders.

## Introduction

Dopamine (DA) is a powerful neuromodulator whose actions impact multiple fundamental behaviors, including novelty and reward salience (1), motivation and attention (2), motor initiation and coordination (3) and executive function (4), among others. Accordingly, altered DA signaling has been implicated in multiple neuropsychiatric disorders such as attention-deficit/hyperactivity disorder (ADHD) (5), autism spectrum disorder (ASD) (6), bipolar disorder (BPD) (7), Parkinson’s disease (8), schizophrenia (9) and substance use disorders (10), though the detailed molecular underpinnings of these conditions remain areas of intense investigation. The presynaptic DA transporter (DAT, *SLC6A3*) tightly regulates the spatiotemporal availability of extracellular DA (11) and aids in replenishing vesicular storages of DA by acting in concert with the *de novo* synthesis machinery (12,13). Abnormal DAT availability in humans with neuropsychiatric disorders has been reported (14,15) and first line treatments for ADHD, i.e., methylphenidate and alpha-methylphenethylamine (d-amphetamine) formulations (16) exhibit high affinity towards DAT (11) and modulate extracellular DA in human subjects (17). Moreover, given abnormal behavior of DAT-manipulated animals (12,18–24) and that the DAT gene *SLC6A3* is implicated in neuropsychiatric diseases (25), ample evidence identifies DAT as a direct contributor to pathological DA dysregulation.

A role of altered DAT function in neuropsychiatric disorders has been advanced by the identification of rare missense variants in the *SLC6A3* gene that encodes DAT in individuals diagnosed with ADHD (26–28), BPD (29) and ASD (30–32). Importantly, these variants have been shown to modulate DAT function (27,28,30,33,34) and/or expression of DAT on the cell surface (27,33,35), and thus are expected to perturb extracellular DA clearance *in vivo*. We and colleagues identified the DAT Ala559Val substitution in two male siblings with ADHD (26), and two unrelated males with ASD (32), a disorder with significant ADHD comorbidity (36–38). Grunhage and coworkers also observed expression of the variant in a girl with BPD (29), a disorder also demonstrating significant familial co-morbidity and shared genetics (39,40). When expressed in a heterologous cell model, DAT Val559 displays normal transporter protein expression, surface abundance and DA uptake rates (33,41). Conspicuously, however, the variant exhibits anomalous DA efflux (ADE) when transfected cells are preloaded with DA (41,42) and increased lateral mobility on the cell surface (43), suggestive of alterations in DAT-associated proteins that influence transporter localization and transport dynamics. DAT dependence of DAT Val559-mediated ADE was established through a block of efflux by methylphenidate and amphetamine, as well as cocaine (41,42). Interestingly, ADE has now been recognized as a functional correlate of multiple disease-associated DAT variants (21,30,34,41,42) and can also be triggered in wildtype (WT) DAT via G protein interactions (44,45), extending the ADE phenotype beyond that seen with rare genetic variants.

*In vivo*, DAT Val559 KI mice (18,46) were found to display normal total DAT expression, as well as DA uptake and tissue DA levels that were indistinguishable from those observed in WT mice (18). In accordance with ADE, however, the striatum of these mice was found to exhibit tonically elevated extracellular DA levels *in vivo* that supports constitutive D2-type autoreceptor (D2AR) activation and blunted vesicular DA release. In keeping with these changes, DAT Val559 mice have been shown to display blunted locomotor responses to methylphenidate and d-amphetamine, heightened locomotor responses to imminent handling, i.e. darting (18), enhanced motivation for reward and waiting impulsivity (19), altered working memory (24) and compulsive reward seeking (23). Together these findings document the value of DAT Val559 mice as a construct-valid model of endogenous alterations in synaptic DA homeostasis that arise from DAT dysfunction. Moreover, the DAT Val559 model provides an *in vivo* platform for the preclinical evaluation of novel therapeutic strategies to ameliorate synaptic DA-associated neurobehavioral disorders, with a particular need for agents without addictive potential.

Various mechanisms have been identified that modulate DAT activity via protein-protein interactions (44,47–49) and post-translational modifications (50,51). DAT is amenable to regulation via presynaptic receptors, either directly or via G proteins, membrane lipids and protein kinases that mediate receptor signaling effects (42,51–54). For example, DA D_2-_type autoreceptors (D2ARs) directly interact with DAT and activation of D2ARs has been shown to promote increased DAT surface trafficking (52,55) via an Extracellular signal-Regulated Kinases 1 and 2 (ERK1/2)-dependent mechanism (52). Notably, as a result of DAT Val559-associated ADE, tonic stimulation of presynaptic D2ARs recruits efflux prone DAT Val559 to the cell surface (56) resulting in a positive feedback-loop that maintains high levels of DAT-dependent ADE. Previously, we demonstrated that D2AR-mediated regulation of DAT in male mice is circuit-specific, as this regulation is seen in nigrostriatal DA projections to the dorsal striatum (DS) but not the mesoaccumbal projections to the ventral striatum (VS) (56). Strikingly, we recently found that the circuit specificity of presynaptic D2AR regulation of DAT is also remarkably sex-specific, with an opposite anatomical pattern of D2AR-DAT coupling seen in female mice, leading to enhanced DAT Val559 surface trafficking in the VS, but not in the DS, a difference that we theorize may contribute to the sex-specific changes observed in multiple DA-linked disorders (24).

Targeting DAT to reduce ADE, as with psychostimulants, invokes use of agents with significant abuse liability (57). Thus, we sought an alternative path for DAT modulation that might normalize DAT Val559 phenotypes with lesser side-effects. In this regard, converging evidence has established a link between kappa opioid receptor (KOR) signaling and functional regulation of DA neurotransmission. For example, KORs are expressed in DA neurons (58) and KOR agonists directly inhibit DA neuron activity (59), which is believed to contribute to stress-related dysphoria (60), resulting from increased secretion of the endogenous KOR ligand dynorphin during stressful events. In addition, exposure to KOR agonists promotes an increase in DAT-mediated DA uptake by increasing DAT surface-trafficking(54). Importantly, KOR antagonism has been found to be well-tolerated in humans (61–65) suggesting opportunities to modulate DA signaling therapeutically via these agents. As a test of this idea, we explored whether KOR signaling can mimic or offset the biochemical and behavioral disruptions observed in DAT Val559 mice. In our current report, using transfected cells and native tissue preparations, we found that activation of KOR increases DAT surface trafficking and DAT N-terminal phosphorylation at threonine 53 (Thr53), a site linked to DAT functional regulation and psychostimulant effects (22,66). Conversely, we found that KOR antagonism reduced basal hyperphosphorylation of DAT Thr53 and surface trafficking observed in male DAT Val559 DS and restored normal rates of activity-dependent vesicular DA release in this model. Finally, KOR antagonism normalized sex-dependent, DAT Val559-induced perturbations in cognitive and locomotor behavior.

## Materials and Methods

Detailed Methods are provided as Supplemental Information (SI)

### Materials

Lipofectamine™ 2000, Dulbecco’s Modified Eagle Medium (DMEM) and other cell culture media were purchased from Invitrogen/Life Technologies, (Grand Island, NY). [^3^H]DA (dihydroxyphenylethylamine [2,5,6,7,8-3H], 63.2 Ci/mmol) and Optiphase Supermix, were purchased from PerkinElmer Inc., (Waltham, MA, USA). U69,593, U0126, protease and phosphatase cocktails were obtained from Sigma-Aldrich (St. Louis, MO). Nor-binaltorphimine dihydrochloride (norBNI) was purchased from Tocris (Bristol, United Kingdom). Reagents for SDS-polyacrylamide gel electrophoresis and Bradford protein assays were from Bio-Rad (Hercules, CA, USA), fetal bovine serum and enhanced chemiluminescence (ECL) reagents were from Thermo Fisher Scientific Inc., (Rockford, IL, USA). Sulfosuccinimidyl-2-[biotinamido] ethyi-1,3-dithiopropionate (EZ link NHS-Sulfo-SS-biotin), Protein A magnetic beads (Dynabeads) and NeutrAvidin Agarose were purchased from Thermo Scientific (Waltham, MA, USA). Anti-Calnexin antibody (Cat# ADI-SPA-860-D, RRID: AB-10616095) was obtained from Enzo Life Sciences, Inc (Farmingdale, NY, USA). QuikChange II XL site-directed mutagenesis kit was obtained from Agilent Technologies (Santa Clara, CA, USA). Peroxidase-affinipure goat anti-rabbit IgG antibody (HRP conjugated secondary antibody (Cat# 111-035-003, RRID: AB-2313567) was acquired from Jackson Immuno Research Laboratories (West Grove, PA). DAT antibody Cat# 431-DATC, RRID: AB-2492076 – was used for experiments using EM4 cells. DAT antibody MAB369 (Millipore Sigma, RRID:AB_2190413) -was used for experiments using acute slices). DAT antibody MABN669 (Millipore Sigma, RRID:AB_2717269) was used for experiments in rat striatal synaptosomes. DAT Thr53 antibody p435-53 (PhosphoSolutions, RRID: AB-2492078) was used to detect DAT phosphorylated at Thr53 in all experiments.

### Cell culture, transfection of DAT cDNAs, and [3H]DA uptake assays

EM4 cells were cultivated as described previously (54). Cells were co-transfected with cDNA constructs encoding myc-KOR plus rDAT-WT, or myc-KOR plus rDAT Ala53, or myc-KOR plus pcDNA3 using Lipofectamine 2000 24 hrs prior to the assay. QuikChange II XL site-directed mutagenesis kit was used to alter rat DAT cDNA sequence encoding threonine at amino acid 53 (Thr53) to encode alanine (Ala53). Uptake of [^3^H]DA into EM4 cells co-transfected with cDNA constructs containing KOR plus rDAT, or rDAT Ala53, or pcDNA3 was assessed following established methods as previously described(54).

### EM4 cell surface biotinylation

Cell surface biotinylation was performed as described previously (54). In brief, transfected EM4 cells were exposed to either vehicle or U69,593 (10 µM) for 30 min at 37°C. Following biotinylation of surface proteins on ice for 30 min, labeled proteins were recovered after cell solubilization using Neutravidin Agaraose resins. Eluates, aliquots of total extracts and unbound fractions were used for SDS-PAGE and immunoblot analysis as described in the SI.

### Animals

All procedures involving animals were approved by the Institutional Animal Care and Use Committees of UT San Antonio Health Sciences Center, Virginia Commonwealth University, or Florida Atlantic University depending on the site of assays, in accordance with the National Institutes of Health Guide for the Care and Use of Laboratory Animals. Male Sprague-Dawley rats (175-350 g body weight) were utilized for *in vivo* chronoamperometry and assays of DAT Thr53 phosphorylation in synaptosomes and tissue. For slice experiments, age-matched 4- to 6-week-old male homozygous mice for either WT or DAT Val559 (genetic background: 75% 129/6 and 25% C57; (18)) were bred from homozygous dams and sires that were derived from heterozygous breeders. Male mice were used for experiments performed in acute slices due to the male bias observed in ADHD diagnosis (67). All biochemical experiments were conducted during the light phase. For behavioral experiments, 6–8-week-old WT and homozygous DAT Val559 littermates derived from heterozygous breeders were utilized. Male mice were used for all behavioral experiments except the novel object recognition test due to the demonstrated sex bias of behavioral phenotypes observed in DAT Val559 mice (24). Behavioral testing was performed during the dark phase of the light cycle under red light. Rodents were maintained in a temperature and humidity-controlled room on a 12:12 h light/dark cycle. Food and water were supplied *ad libitum*. All efforts and care were taken to minimize animal suffering and to reduce the number of animals used. As alternatives to brain tissues, cell culture models were utilized.

### Treatment of striatal synaptosomes with U69,593 ± U0126

Male Sprague-Dawley rats (175–300 g) were decapitated, and striata were rapidly dissected, weighed, and placed in ice-cold sucrose phosphate (SP) buffer (0.32 M sucrose and 10 mm sodium phosphate, pH 7.4). Tissues were homogenized in ice-cold SP buffer with 15 strokes in a glass/Teflon homogenizer and centrifuged at 3000 × *g* for 3 min at 4 °C. Supernatant fractions were re-centrifuged at 17,000 × *g* for 12 min, and the resulting P2 pellet enriched for synaptosomes was resuspended to 20 mg/mL original wet weight in ice-cold SP buffer. Synaptosomes were treated with vehicle or 50 µM U0126 for 15 min at 30°C, followed by treatment with vehicle or 10 µM U69,593 for an additional 15 min at 30°C. After treatment, samples were subjected to SDS-PAGE followed by immunoblotting for total DAT and DAT phosphorylated at Thr53).

### p-Thr53 DAT levels from rats injected with vehicle or U69,593

Male rats were injected s.c. with vehicle or 0.32 mg/kg U69,593 and sacrificed at the indicated times noted in Figure 3. DS or VS were dissected, weighed and kept frozen until analyzed. The pre-weighed samples were homogenized with a Polytron PT1200 homogenizer (Kinematica, Basel, Switzerland) for 8 s in ice-cold SP buffer, and centrifuged at 3000 *× g* for 3 min at 4 °C. Supernatant fractions were re-centrifuged at 17,000 *× g* for 12 min to obtain a membrane pellet. The pellet was resuspended to 20 mg/mL original wet weight in ice-cold SP buffer. Samples were subjected in duplicate to SDS-PAGE and western blot for total and p-Thr53 DAT, and p-Thr53 DAT staining quantified by normalization to the time-matched controls.

### Analysis of in vivo DA clearance in NAc

High-Speed *in vivo* chronoamperometry, conducted as previously described (68), was used to determine the effect of KOR modulators on DA clearance in NAc of anesthetized rats. Briefly, carbon fiber electrodes were attached to multi-barrel glass micropipettes. Rats were anesthetized with chloralose (85 mg/kg, i.p.) and urethane (850 mg/kg, i.p). Electrodes were lowered into the NAc (69)using a stereotaxic frame. Individual micropipette barrels were filled with DA (200 µM), U69,593 (885.7 µM, barrel concentration), norBNI (400 µM, barrel concentration) and vehicle and ejected (100–150 nL) using a Picospritzer II (General Valve Corporation, Fairfield, NJ)(68). DA was pressure-ejected until a stable baseline signal was established (typically after 3-4 ejections). The effect of U69,593, norBNI, and vehicle on DA signals was quantified 2 min after a stable baseline DA signal was established. See SI for experimental details.

### Surface biotinylation and immunoprecipitation of p-Thr53 DAT using acute mouse brain slices (56)

#### Acute slice preparation

300 µm thick acute coronal slices were prepared from 4-6 week-old male mice. Following recovery, plane-matched hemi slices were exposed to either vehicle or drug (U69,593: 10 µM for7 min. norBNI: 1 µM for 20 min) at 37 °C. After drug treatments, the slices were subjected to three rapid washes and two 5-min washes on ice. For immunoprecipitation studies, VS and DS were dissected at this point and tissue was flash frozen in liquid nitrogen and stored at -80 °C until processed. For biotinylation assays, the slices were exposed to 1 mg/mL EZ link NHS-Sulfo-SS-biotin for 30 min on ice with constant oxygenation. Following washes, tissue was solubilized in RIPA buffer and biotinylated preoteins were recovered using Streptavidin Agarose beads. The experimental procedures for the biotinylation and immunoprecipitation assays are described in detail in (56) and in the SI.

### Behavioral testing

Y Maze, Open Field test, cocaine-induced locomotor activation, and the novel object recognition (NOR) test were performed using 6-8 week old male and female (only for the NOR test) mice in the Stiles-Nicholson Brain Institute Neurobehavior Core of Florida Atlantic University based on established protocols (24) and are described in detail in the SI.

### In vivo microdialysis

Microdialysis was performed using 6–8-week-old male mice. Surgeries to insert microdialysis guide cannulae were performed as described earlier (18,69). A 5 mm guide cannula was lowered into the DS and affixed to the skull with dental cement, supported by three 1.6 mm screw. Mice were allowed to recover for 6 days. 12 hr prior to the experiment a microdialysis probe with an active membrane length of 3 mm was inserted into the guide cannula and microdialysis, sample collection and analsyis were performed as described in (18). After collection of the three baseline fractions, mice received an i.p. injection of norBNI (10 mg/kg) and cocaine hydrochloride (10 mg/kg; i.p.) was injected 80 min thereafter. Changes in extracellular DA were expressed as fold increase of basal DA (i.e. the average DA of the first three basal dialysates).

### Statistical analyses

Prism 7 (GraphPad, San Diego, CA) was used for data analysis and graph preparation. Values are presented as mean ± standard deviation (SD). One-way or two-way ANOVA were used followed by post hoc testing for multiple comparisons. The type and results of post hoc tests are noted in the Figure legends. Two-tailed, unpaired Student’s t-tests were performed for comparisons between two groups. P values ≤ 0.05 were considered statistically significant.

### Data Availability

All data will be provided by the corresponding authors upon request.

## Results

### KOR regulation of DAT activity requires the capacity to phosphorylate DAT at N-terminal residue Thr53

To initiate our evaluation of KOR manipulation as a strategy for offsetting DAT-dependent alterations in DA homeostasis, we transiently transfected EM4 cells with rat cDNAs encoding KOR and DAT (rKOR and rDAT, respectively). In agreement with a previous study (54), activation of KOR enhanced DAT-mediated uptake in a time and concentration dependent manner (Figure 1 A, B). We selected 10 µM U69,593 and a 15 min pre-treatment for subsequent *in vitro* experiments and found that exposure to U69,593 significantly increased DA transport V_max_ (336.1 ± 18.12 (SD) versus 551.2 ± 21.79 (SD) pmol/min/10^6^ cells), whereas DA K_M_ was unaffected (Figure 1C).

**Figure 1:**
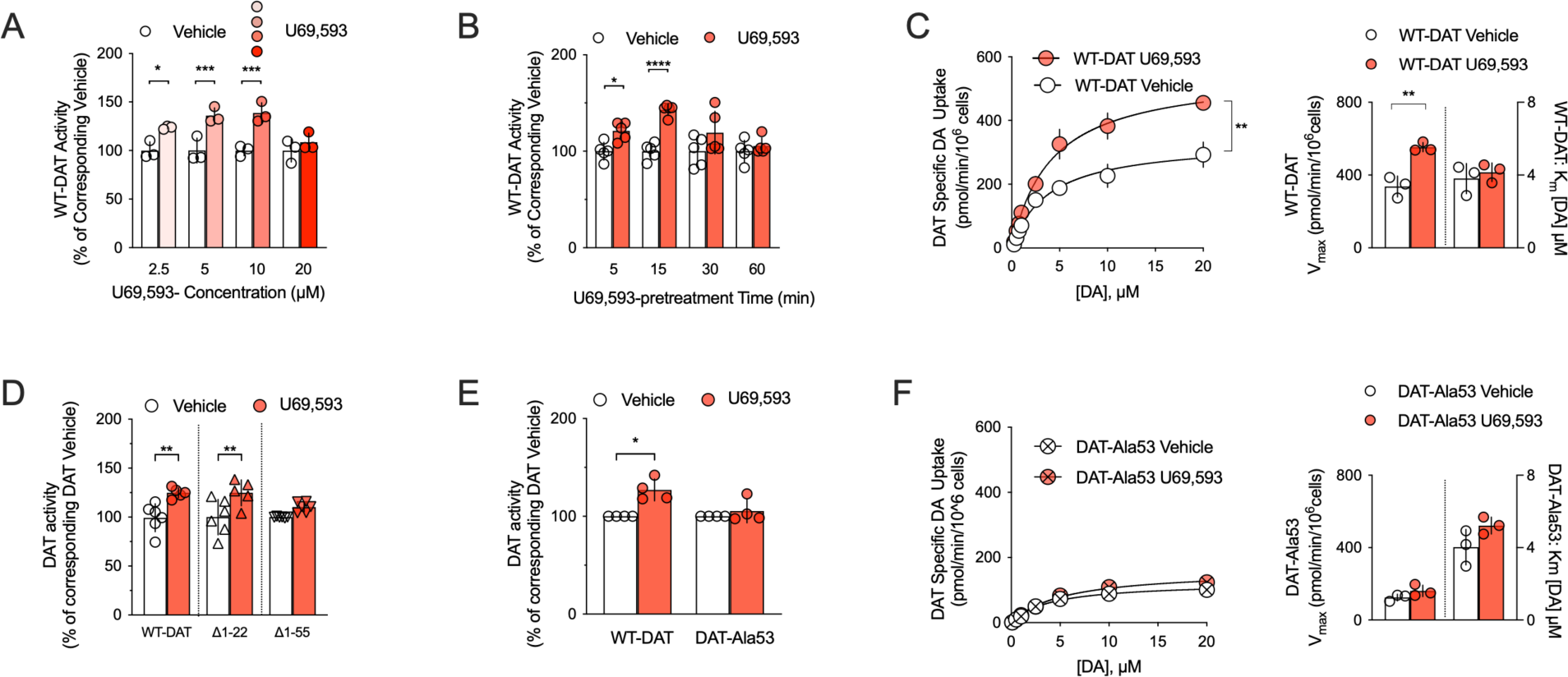
The KOR agonist U69,593 increases DAT-mediated uptake via a Thr53 dependent mechanism. EM4 cells were transfected with DAT and KOR and uptake of [^3^H]DA was assessed as described in Methods. **A**: U69,593 (10 µM) increased WT DAT-mediated uptake in rDAT/rKOR cotransfected EM4 cells in a concentration-dependent manner as compared to vehicle (n= 3 independent observations per condition, one-way ANOVA, followed by Bonferroni post-hoc tests). **B**: Treatment with 10 µM U69,593 enhanced WT DAT specific uptake in a time-dependent fashion (n= 5 independent observations per condition, one-way ANOVA, followed by Bonferroni post-hoc tests). **C**: Incubation with 10 µM U69,593 increased WT DAT V_max_ (two-tailed unpaired t-test) without effect on K_M_ (n=3 independent observations per condition, two-tailed Student’s unpaired t-test). **D**: Treatment with U69,593 significantly augmented WT DAT-mediated uptake when compared to vehicle in cells that expressed KOR plus either WT DAT or a truncated version of DAT, lacking the first 22 N-terminal residues (DAT Δ1-22). In contrast, truncation of the first 55 N-terminal residues (DAT Δ1-55 DAT) prevented KOR-agonist induced increases in [^3^H]DA uptake. (n=5-6 independent observations per condition, one-way ANOVA, followed by Bonferroni post-hoc tests) **E**: Site-directed mutagenesis of DAT Thr53 to alanine (DAT-Ala53) rendered transporter-mediated DA uptake insensitive to pre-treatment with U69,593 at 5 and 10 µM (n=4 observations per condition, One-way ANOVA, followed by Bonferroni posthoc tests). **F**: U69,593 did not affect the kinetic parameters K_M_ and V_max_ of the DAT-Ala53 mutant (n=3 independent observations per condition, Student’s two-tailed unpaired t-test) Data are shown as the mean with error bars reflecting the SD *=P<0.05; **=P<0.01; ***=P<0.001; ****=P<0.0001.

The site(s) or motifs in DAT through which KOR regulates DAT function have not been defined. Comprehensive research has identified DAT phosphorylation as a key mechanism that determines DAT activity and membrane trafficking (51,70,71). We focused on the N-terminus due to evidence that the MAP kinase ERK1/2 supports KOR regulation of DAT (54) and a conserved, N-terminal proline-directed kinase consensus phosphorylation motif containing Thr53 that can be phosphorylated in *vitro* by ERK1 (72) has been implicated in DAT function *in vitro* and *in vivo* (66,73,74). We first generated two truncated versions of rDAT, lacking either the first 22 (Δ1-22) or 55 (Δ1-55) N-terminal residues. Despite the loss of multiple phosphorylation sites that are found within the most distal N-terminus (51,71), rDAT Δ1-22 remained amenable to regulation via activation of rKOR, as treatment with U69,593 significantly increased DA uptake in EM4 cells coexpressing rDAT ΔN1-22 and rKOR (Figure 1D). In contrast, no uptake stimulation by U69,593 was evident in cells transfected with rKOR and rDAT ΔN1-55. Given that Thr53 is located in the region that distinguishes ΔN1-55 from Δ1-22, we replaced Thr53 with an alanine (rDAT Ala53) to preclude phosphorylation at this site. Single point and saturation uptake analyses in EM4 cells transfected with rKOR and either WT rDAT or rDAT Ala53 revealed an absolute requirement for Thr53 phosphorylation potential for U69,593 to increase DA uptake (Figure 1E-F). Saturation analysis also demonstrated that a reduction in V_max_ supports the basal reduction in rDAT Ala53 DA uptake (124.4 ± 18.45 (SD) pmol/min/10^6^cells)) evident in comparison to WT rDAT (337.4 ± 57.65 (SD) pmol/min/10^6^cells) (Figures 1C, 1F).

### KOR agonism increases rDAT surface levels in a Thr53-dependent manner in EM4 cells

Given the well-established relationship between DAT surface expression and DA uptake, we next sought to determine whether the Thr53-dependent KOR elevation of rDAT-mediated uptake correlates with an increase in transporter surface expression. To pursue this objective, we biotinylated transfected cells with the membrane-impermeant reagent EZ link NHS-Sulfo-SS-biotin and recovered biotinylated surface proteins from total protein extracts. Exposure of cells to U69,593 had no effect on total levels of either WT rDAT or rDAT Ala53 (Figure 2A). Western blots of total extracts revealed WT rDAT to migrate as distinct bands of ∼55-60 kDa and ∼85-90 kDa, typical of rDAT transiently transfected cells and linked to different transporter species bearing mature versus immature states of N-glycosylation (75). In contrast to the WT rDAT protein profile, extracts from cells expressing the DAT Ala53 mutant were depleted of the 85-90 kDA species. Analysis of rDAT species in biotinylated samples showed that U69,593 significantly increased WT rDAT surface density for both the ∼55-60 kDa and ∼85-90 kDa DAT bands. In contrast, neither species of rDAT Ala53 was affected by U69,593 treatment (Figure 2B). To detect rDAT Thr53 phosphorylation (p-Thr53), we probed blots with a DAT p-Thr53 specific antibody (66). We found that U69,593 treatment significantly increased immunoreactivity for the ∼85-90 kDa isoform of DAT p-Thr53 labelled proteins in total extracts from WT rDAT transfected cells, whereas increased DAT p-Thr53 labelling of the WT rDAT ∼55-60 kDa isoform was not detected. Importantly, no DAT-specific bands of either isoform were detected in surface-samples from cells transfected with the rDAT Ala53 variant (Figure 2C). Finally, when we analysed blots from extracts recovered after surface biotinylation, we detected a significant increase in DAT p-Thr53 immunoreactivity for both isoforms of WT DAT, whereas no DAT p-Thr53-specific bands were detected in the case of the Ala53 variant (Figure 2D). Of note, the DAT p-Thr53 antibody detected some non-specific bands, which were also present in mock-transfected cells (* in Figures 2C and D) and therefore not considered for analysis. Together, our findings indicate that capacity for Thr53 phosphorylation is essential for enhanced surface trafficking of rDAT in response to rKOR agonism in EM4 cells.

**Figure 2:**
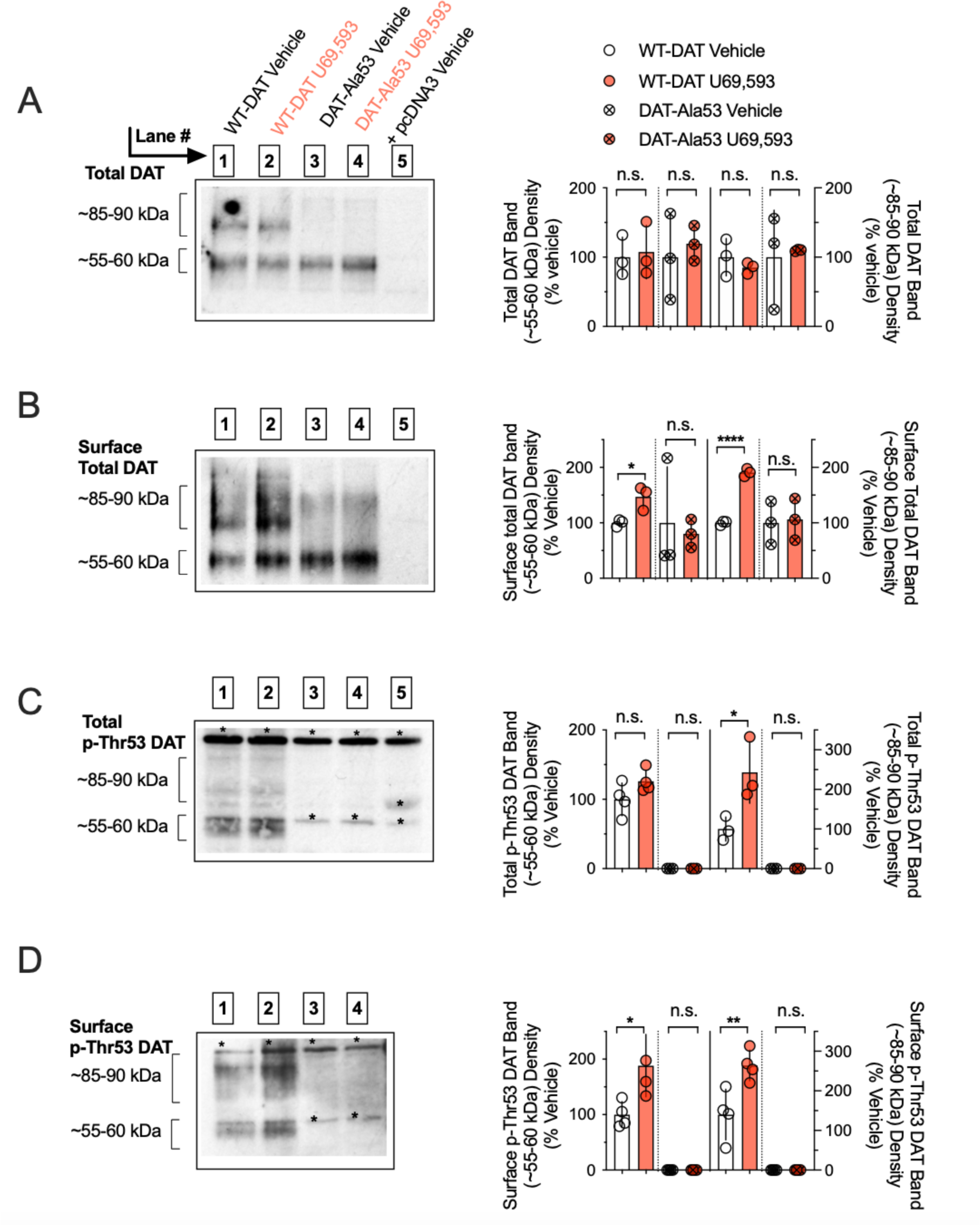
Capacity to phosphorylate rDAT at Thr53 is required for rKOR activation to elevate transporter surface expression. EM4 cells were transfected with either WT DAT or DAT Ala53 plus rKOR, subjected to U69,593 and biotinylated as described in Methods. **A**: Treatment with U69,593 did not affect WT DAT band densities at ∼55-60 kDa and ∼85-90 kDa, derived from total protein lysates (n=3 independent observations per condition, Student’s two-tailed unpaired t-test). **B**: Incubation with U69,593 increased the WT DAT band densities at ∼55-60 kDa and ∼85-90 kDa, derived from biotinylated protein lysates (unpaired, two-tailed t-test). No effect was observed on the DAT Ala53 band densities at ∼55-60 kDa (n=3 independent observations per condition, Student’s two-tailed unpaired t-test). C and D: Protein lysates were immunoblotted with p-Thr53 antibody. Bands marked with an asterisk indicate non-specific immunoreactivity. **C**: In total protein lysates, incubation with U69,593 increased the density of the p-Thr53 immunoreactive band at ∼55-60 kDa for WT DAT (Student’s unpaired, two-tailed t-test). No effect was observed on the band densities at ∼85-90 kDa. No specific bands were detected for the DAT Ala53 variant (n=3-5 independent observations per condition, Student’s two-tailed unpaired t-test). **D**: U69,593 augmented band densities at ∼55-60 and ∼85-90 kDa for biotinylated WT DAT (n=5 independent observations per condition, Student’s two-tailed unpaired t-test). No specific bands were detected for the DAT Ala53 variant. All bars represent the mean with error bars reflecting the SD. Individual quantifications are displayed as individual symbols. *=P<0.05; **=P<0.01; ****=P<0.0001. n.s.=not significant

### Pharmacological activation of KOR increases rDAT phosphorylation at Thr53 in striatal synaptosomes and enhances DA clearance in vivo

Having established in transfected cells that a capacity for rDAT Thr53 phosphorylation plays an essential role in KOR-dependent elevations of transporter surface trafficking and DA uptake, we next sought to determine whether these findings could be replicated in native tissue. As shown in Figure 3A, treatment of rat striatal synaptosomes with U69,593 promoted a significant increase in p-Thr53 DAT immunoreactivity that could be blocked by inhibition of mitogen-activated protein kinase kinase (MEK1/2) using U0216, which lies upstream of ERK 1/2 (Figure 3A). To assess whether KOR activation can elevate p-Thr53 DAT *in vivo*, male rats were injected with U69,593 (0.32 mg/kg, i.p.) or vehicle and sacrificed at the time points shown in Figure 3B and C. To parse out whether circuit-specific effects that shape the outcome of D2AR regulation of DAT surface trafficking (24,56) also exist for KOR, we analysed the impact of systemic U69,593 on p-Thr53 DAT by western blot in both the DS (Figure 3B) and VS (Figure 3C). For both DS and VS, systemic U69,593 caused a time-dependent increase in p-Thr53 DAT that peaked at 120min post injection. Considering the effects of KOR activation to increase DAT-mediated DA uptake in cells (Figure 1 and (54)), we sought *in vivo* evidence for increased DAT function following KOR activation via assessment of DA clearance time using high-speed *in vivo* chronoamperometry (68,76). A carbon fiber electrode-micropipette assembly was lowered into the NAc (Figure 3D) permitting DA to be pressure ejected, followed by recording of DA levels as a function of time. Representative traces depicting DA clearance before and after administration of U69,593 are shown in Figure 3E. We found that local injection of U69,593 decreased DA clearance time compared to predrug baseline, quantified as the time required to clear 50% (T50) and 80 % (T80) of the DA-induced amperometric signal (Figure 3F). In contrast, pre-treatment with the KOR antagonist norBNI significantly increased clearance time consistent with a non-saturating level of endogenous KOR activation under our recording conditions (Suppl. Figure 2).

**Figure 3:**
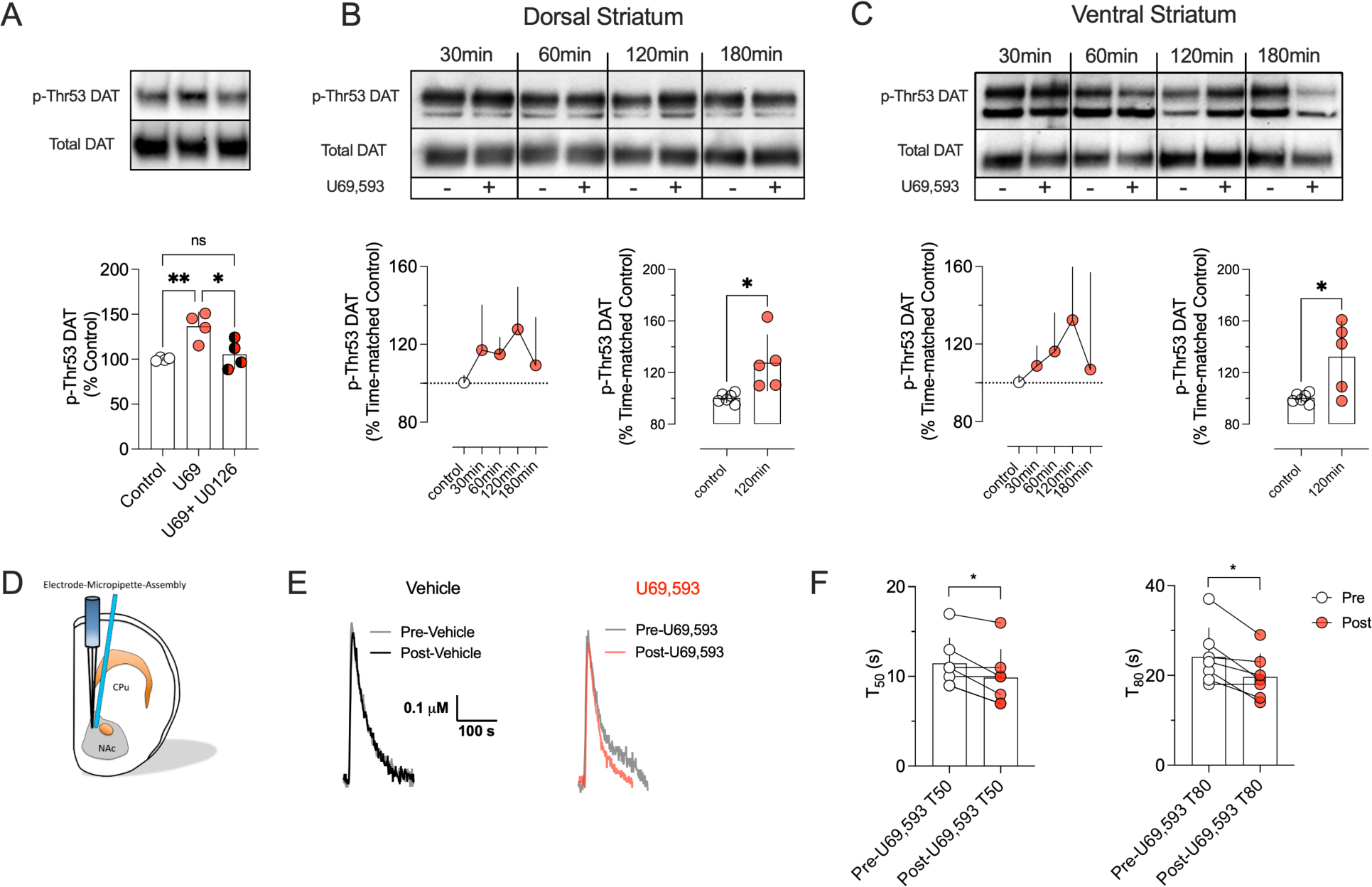
KOR-agonism enhances phosphorylation of DAT at Thr53 in native preparations and augments DAT-mediated clearance in the NAc *in vivo*. **A)** Male rat striatal synaptosomes were treated with vehicle or 50 µM U0126 for 15 min at 30°C, followed by treatment with vehicle or 10 µM U69,593 for an additional 15 min 30°C. After treatment, samples were subjected to SDS-PAGE and western blotted for total and p-Thr53 DAT. Upper panel shows representative western blots. Lower panel shows quantification of p-Thr53 DAT staining as a percentage of basal levels. (n=4; one-way ANOVA, Tukey’s multiple comparisons test). **B and C)** Rats were injected s.c. with vehicle or U69,593 (0.32 mg/kg) and sacrificed at indicated times. Tissue from the dorsal **(B)** or ventral striatum **(C)** were western blotted for total and p-Thr53 DAT in duplicate. Signals from the U69,593 treated samples were compared to the time-matched control. The upper panels show representative western blots for the dorsal and ventral striatum with each time-matched control followed by U69,593 treatment. The lower panels show quantification of the p-Thr53 DAT staining as a percentage of the time-matched control (Two-tailed, unpaired t-test). **D)** Schematic representation of the electrode-micropipette assembly that was lowered into the NAc of anaesthetized rats to allow for *in vivo* chromatographic measurement of DA clearance rates. **E)** Representative oxidation currents converted to micromolar values observed upon pressure ejection of DA before (gray traces) and 2 min after intra-NAc injection of vehicle (black trace), U69,593 (890 µM, barrel concentration, red trace). The leftward-shift in the representative trace following U69,593 injection is indicative of increased DA clearance. **F)** U69,593 decreased the clearance time of exogenously applied DA in the NAc (T_50_ and T_80_; n= 7 observations per condition, two-tailed paired t-test) when compared to the pre-drug value. Bars indicate the mean and error reflect SD. Individual values from each animal are reflected by the corresponding symbols. *denotes P<0.05; **denotes P<0.01, ns = not significant.

### Targeting KOR to normalize aberrant surface trafficking and phosphorylation of the disease associated DAT variant Val559 in vivo

DAT Val559 expression in male mice is associated with increased surface trafficking of DAT and elevated levels of p-Thr53 DAT in the DS but not the VS (56). The ability of KORs to regulate DAT via phosphorylation of Thr53 raised the possibility that pharmacological antagonism of mKORs might normalize the tonically elevated surface expression of efflux-prone mDAT Val559 proteins. To determine whether mDAT Val559 remains amenable to regulation via mKOR in native brain preparations, we first treated acute coronal slices containing either DS or VS of WT mDAT or mDAT Val559 *in vitro* with U69,593 (10 µM, 7 min), extracting DAT proteins thereafter for analysis of transporter surface trafficking and p-Thr53 levels via western blots. *In vitro* activation of mKOR increased surface trafficking of DAT independent of genotype in both the DS (Figure 4A) and VS (Figure 4C). As with surface transporter density, and as previously reported (56), blots of mDAT Val559 extracts from male DS (Figure 4B) but not VS (Figure 4D), demonstrated significantly elevated levels of p-Thr53 DAT relative to WT mDAT under vehicle-treated conditions. Whereas U69,593 (10 µM, 7 min) increased levels of p-Thr53 DAT in extracts both from DS (Figure 4B) and VS (Figure 4D) of WT male mice as well as VS from DAT Val559 male mice (Figure 4 D), this effect was blunted in male DAT Val559 preparations from the DS (Figure 4).

**Figure 4:**
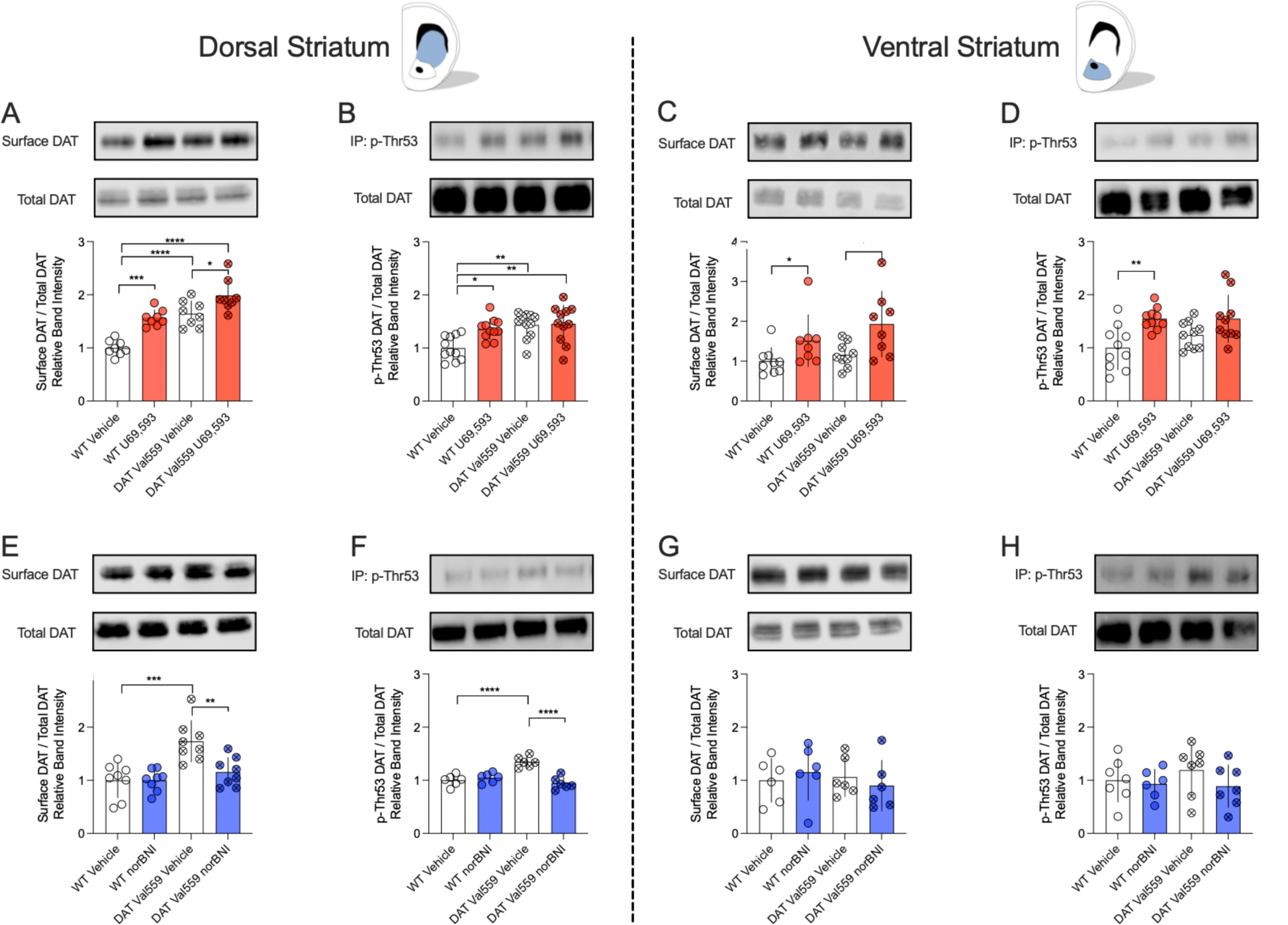
mKOR agonism induces enhanced mDAT surface expression and phosphorylation at mDAT Thr53 while mKOR antagonism normalizes enhanced surface expression and Thr53 phosphorylation of mDAT Val559 in acute brain slices containing the DS or VS. **A-D)** Acute slices containing the DS and/or the VS were exposed to 10 µM of U69,693 or vehicle for 7 min and surface expression and phosphorylation of p-Thr53 DAT was assessed. **A** and **B)** In the DS, treatment with U69,593 increased surface DAT levels (A) (n=9; two-way ANOVA, Šídák’s multiple comparisons test) as well as phosphorylation of mDAT at Thr53 (B) (n=10; two-way ANOVA, Šídák’s multiple comparisons test). Higher basal phosphorylation levels at Thr53 were detected for the DAT Val559 when compared to WT DAT in the DS (n=11; two-way ANOVA, Šídák’s multiple comparisons test). **C)** In the VS, KOR-agonist treatment increased surface mDAT levels independent of genotype as compared to vehicle treatment (n=8; two-way ANOVA, Šídák’s multiple comparisons test). **D)** Treatment with U69,593 enhanced WT mDAT phosphorylation at Thr53, but remained without effect on DAT Val559 phosphorylation. **E-H)** Acute coronal slices containing the DS or the VS were treated with the KOR antagonist norBNI (1 µM) for 20 min and DAT surface expression and phosphorylation at Thr53 were determined. **E** and **F)** In the DS, antagonism of mKOR reduced the enhanced surface expression **(E)** (n=8; two-way ANOVA, Šídák’s multiple comparisons test) and Thr-53 phosphorylation **(F)** (n=6; two-way ANOVA, Šídák’s multiple comparisons test) of tansporters in DAT Val559 mice. No effect of norBNI was detected on WT DAT. **G** and **H)** Treatment with norBNI remained without effect on DAT surface expression **(G)** (n=6, two-way ANOVA, Šídák’s multiple comparisons test) and Thr-53 phosphorylation **(H)** (n= 7, two-way ANOVA, Šídák’s multiple comparisons test) in the VS. All bars show the mean and SD. Symbols reflect individual observations. * = *P* ≤ 0.05, ** = *P* ≤ 0.01, *** = *P* ≤ 0.001, **** = *P* ≤ 0.0001

We found that norBNI (1 µM, 20 min) produced no effect on either surface density or DAT p-Thr53 levels when applied to WT DAT DS slices. However, treatment with norBNI in vitro normalized the elevations observed in both transporter surface density (Figure 4E) and p-Thr53 DAT (Figure 4F) in the DS of DAT Val559 mice relative to WT controls. In the VS, norBNI was without effect on transporter trafficking or p-Thr53 levels of either WT or DAT Val559 mice (Figure 4 G and H). To establish that this effect is of relevance in vivo, we treated pairs of WT and DAT Val559 mice either with saline or norBNI (10 mg/kg, i.p.) 30 min prior to the collection of acute coronal slices, which were then subjected to immediate surface biotinylation. Enhanced surface expression of mDAT Val559 relative to WT mDAT was evident in slices prepared from saline treated animals, whereas no difference in surface density was observed following norBNI treatment (Supplementary Fig 3).

### In vivo KOR antagonism rescues vesicular DA release and behavioral deficits in DAT Val559 mice

Consistent with tonic D2AR activation and D2AR-mediated suppression of vesicular DA release, *ex vivo* studies with striatal slices from male DAT Val559 mice revealed a significant reduction of evoked vesicular release of DA (18). Additionally, *in vivo* microdialysis in the DS of these animals revealed a loss of cocaine (10 mg/kg, i.p.) induced elevations in extracellular DA (20). We hypothesized that mKOR inhibition, due to its ability to reduce the elevated basal surface trafficking of efflux-prone mDAT Val559 should restore vesicular DA release in vivo. Indeed, as assed by in vivo microdialysis, when male mice were treated with norBNI (10 mg/kg; i.p.) 80 min prior to an i.p. injection of cocaine (10 mg/kg), we found that cocaine now evoked a robust increase in extracellular DA in DAT Val559 mice, equivalent to that seen with WT mice (Supplementary Figure 4 A-C).

To determine whether a restoration of vesicular DA release might lead to a normalization of behaviors found to be disrupted in male and female DAT Val559 mice. First, we chose the Y-maze test of spontaneous alternation, a measure whose reduction is considered a symptom of disrupted working memory or compulsive repetitive behavior. Male but not female DAT Val559 mice display a robust reduction in alternations in the Y maze, when compared to WT littermates, driven by immediate return to a previously visited arm (direct revisits) (Figure 5A through D and (24)) that was not explained by changes in locomotor activity (Figure 5C and (24)). Strikingly, an acute injection of norBNI (10 mg/kg, i.p. 30 min prior to initial Y maze testing) normalized the alternation deficit of DAT Val559 mice (Figure 5B) as well as the direct revisits monitored in the task (Figure 5D) without an effect on locomotor behavior (Figure 5 C).

**Figure 5:**
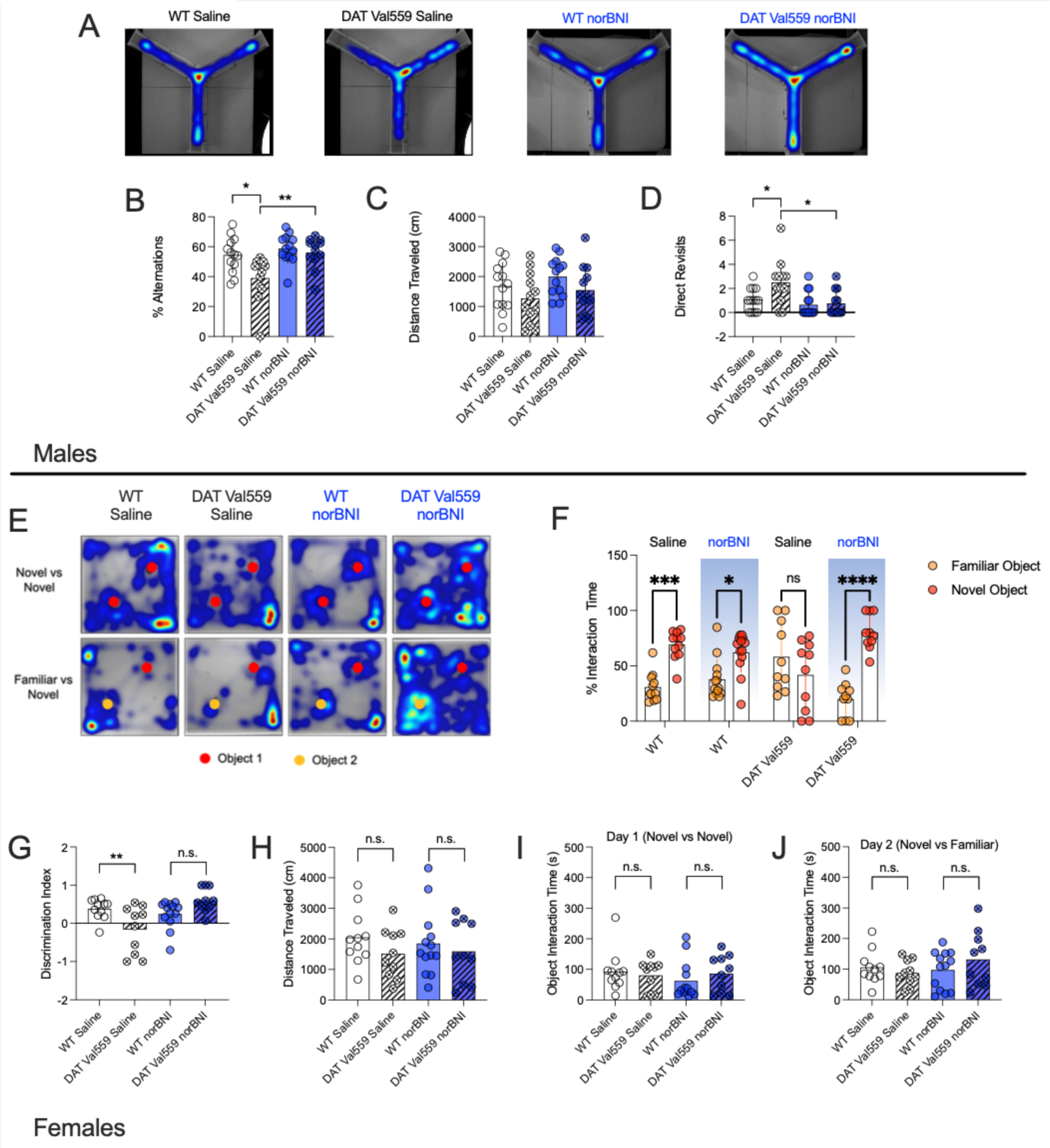
Treatment with norBNI normalizes sex-specific behavioral phenotypes of male and female DAT Val559 mice. A-D) Male homozygous DAT Val559 mice and their WT littermates were injected with saline (vehicle; n= 13 for WT and 12 for DAT Val559, respectively) or norBNI (10 mg / kg, i.p.; n= 13 for WT and 12 for DAT Val559, respectively) 30 min prior to testing, placed into the center of the open Y-maze and the number of alternating arm entries, distance travelled and direct revisits were recorded as described in Methods. A) Representative heat maps showing the explorative behavior of WT and Val559 mice following vehicle or norBNI administration. Time spent in each area is directly correlated to the color gradient ranging from dark blue to dark red, with the latter indicating highest value. B) Systemic administration of norBNI 30 min prior to the test normalized the deficit in the percentage of alternations of DAT Val559 mice when compared to WT control mice. C) No acute effect of norBNI was observed for total distance travelled during the test session. D) Administration of norBNI reduced the number of direct revisits of DAT Val559 mice. E-J) The NOR task was performed with saline treated female WT (n=11) and DAT Val559 littermates (n=10) and compared to norBNI treated (10 mg/kg, i.p., 30 min prior to testing) female WT (n=13) and littermate DAT Val559 (n=10) mice. E) provides representative heat maps representing location in relation to objects (circles) used in NOR sessions. The relative interaction time with the familiar versus the novel object and the discrimination index are shown in (F) and (G), respectively. H) shows the total distance travelled and the total object interaction times on day 1 (two novel objects) and day 2 (one novel and one familiar object) are shown in panels (I) and (J), respectively. Data are given as mean and SD and were analyzed using two-way ANOVA with Šídák’s multiple comparisons test. *=P<0.05, **=P<0.01, ***=P<0.001, ****=P<0.0001, n.s.= not significant.

DAT Val559 affects DA circuits in a sex-dependent manner that leads to distinct behavioral outcomes in males versus females (24). To determine if systemic treatment with norBNI (10 mg/kg, i.p.) is also capable of reversing a female-specific phenotype of DAT Val559 expression, we implemented the novel object recognition (NOR) test, where female but not male DAT Val559 mice display strong differences in novel object preference and discrimination, as female DAT Val559 mice do not explore a novel object to the same extent as WT controls (24). Again, treatment with norBNI (10 mg/kg, i.p.) restored the discrimination index and time spent with the novel object of female DAT Val559 mice to levels that were comparable to those observed in female WT mice (Figure 5E through J). Importantly, deficits in the relative interaction time and discrimination index are not confounded by locomotor activity or total object interaction time on testing day 1 or testing day 2 (Figure 5 F,G,H,I and J, respectively), as all these measures remained unchanged in female DAT Val559 mice. Consequently, these findings demonstrate that mKOR antagonism can normalize behavioral changes unique to either male or female DAT Val559 mice.

NorBNI is associated with long lasting effects *in vivo* (77,78). Therefore, we reassessed Y maze activity in the same male experimental subjects as reported above, one week after the initial test. One week following the initial study, the vehicle treated male DAT Val559 mice continued to display a deficit in alternations (Supplementary Figure 5B) that was not driven by altered locomotor activity (Supplementary Figure 5C) but rather the result of an increase in direct revisits when compared to WT controls (Supplementary Figure 5D).In contrast, the norBNI treated DAT Val559 mice displayed a significantly altered alternation pattern when compared to vehicle treated DAT Val559 mice (Supplementary Figure 5B). Moreover, the norBNI treated DAT Val559 group exhibited a direct revisit pattern that was statistically indistinguishable from vehicle treated WT controls (Supplementary Figure 5D), whereas no drug or genotype effect on total distance travelled during the Y maze test sessions was observed one week post norBNI treatment (Supplementary Figure 5C).

We further examined the behavior of the vehicle and norBNI treated cohorts one week post administration in the open field test (OFT). Unlike DAT KO mice (12), male (or female) DAT Val559 mice are not spontaneously hyperactive, though male DAT Val559 mice display increased locomotor reactivity (darting) in response to imminent handling (18). In the OFT, male DAT Val559 mice displayed locomotor suppression (distance travelled, rearing and stereotypy), as compared to WT littermates (Figure 6A,B). Remarkably, injection of norBNI 7 days prior to testing normalized these measures (Figure 6D,E). We hypothesize that the reduced locomotion of the saline treated DAT Val559 mice may represent heightened anxiety post handling, since DAT Val559 mice demonstrated reduced center time in the open field (Fig 6C), a behavioral measure known to be responsive to anxiolytics (79). Systemic norBNI also normalized this measure (Figure 6C). DAT Val559 further mice display a blunted locomotor response to psychostimulants, including cocaine (10 mg/kg, i.p.) (18,20). Pretreatment with norBNI (10 mg/kg, i.p.) one week prior to testing restored the locomotor response of male DAT Val559 mice to systemic cocaine. In the norBNI treated group, the increase in cocaine-induced locomotor activity was concomitant with an increase in time spent in the center of the chamber, with the latter measure being absent in saline treated controls (Figure 6 F and G).

**Figure 6.**
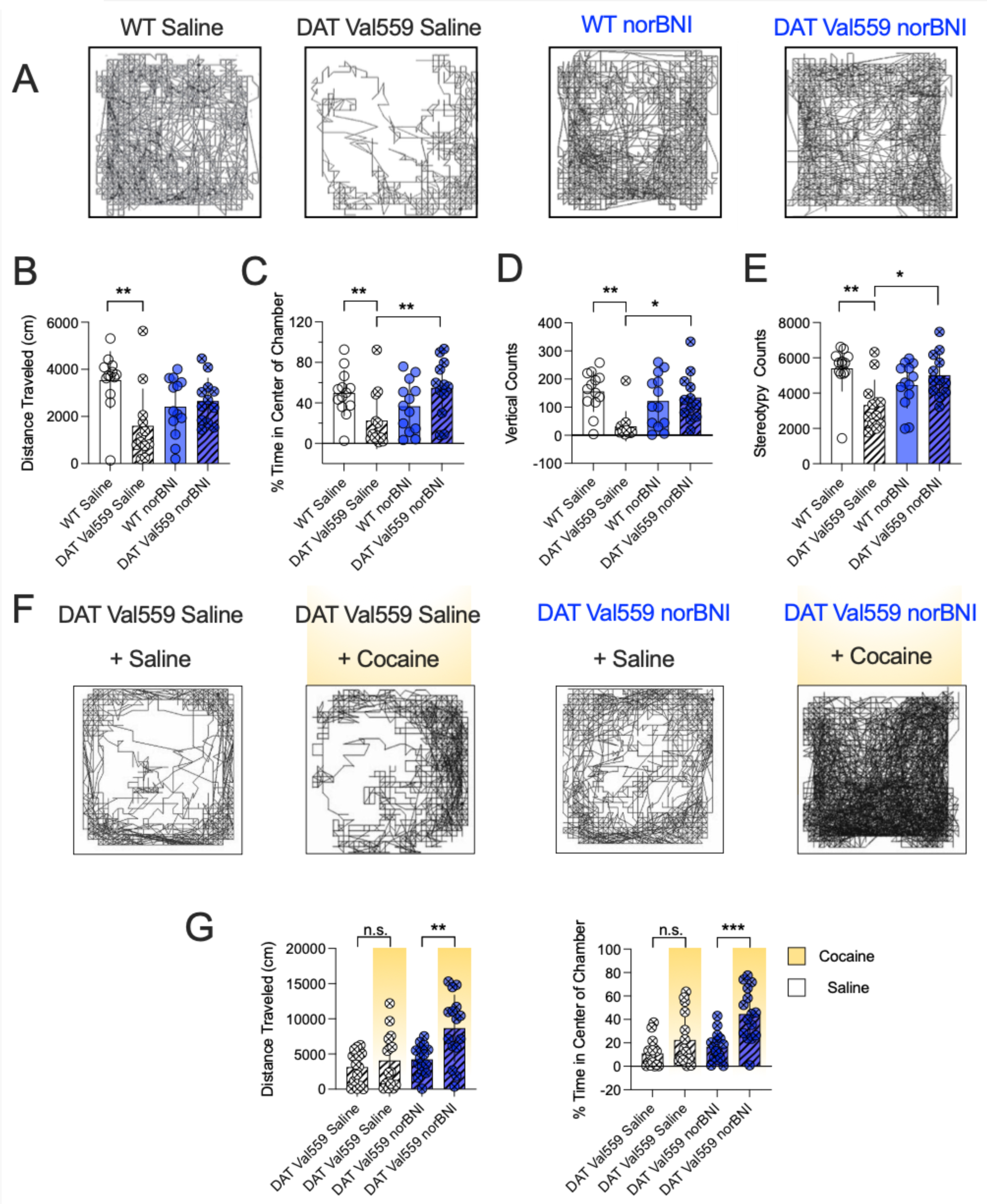
norBNI treatment of male DAT Val559 mice normalizes abnormal locomotor responses to injection stress and locomotor response to systemic cocaine as monitored in the open field test. A-E) Homozygous male DAT Val559 and WT littermates were injected with saline (vehicle) or norBNI (10 mg/kg, i.p.) and placed in open activity chambers one week post injection. A) Representative activity traces of WT and DAT Val559 mice. DAT Val559 mice previously injected with vehicle (saline) displayed significantly less B) forward locomotor activity C) spent less time in the center of the activity chamber and exhibited fewer D) vertical counts and E) stereotypies when compared to WT mice. No differences between genotypes were detected when the mice were pretreated with norBNI. F-G) Homozygous male DAT Val559 were injected with saline or cocaine (10 mg/kg, i.p.) one week post administration of norBNI (10 mg/kg, i.p.) and placed into open activity chambers. F) Representative traces of male DAT Val559 mice injected with the indicated drug combinations. G) Total distance travelled and time spent in the center of the chamber of male DAT Val559 mice injected with the indicated combinations of saline, norBNI and cocaine. All bars show the mean and SD. n=12-13 individual animals per group, data in B, C, D, E and G were analyzed with two-way ANOVA, Šídák’s multiple comparisons test. Symbols reflect individual animals. * = *P* ≤ 0.05, ** = *P* ≤ 0.01, *** = *P* ≤ 0.001, ns = not significant.

## Discussion

Dysfunction in DAergic signaling has been linked to multiple neuropsychiatric disorders including ADHD (80), schizophrenia (81) and ASD (82). Initial efforts to model disturbed DA-signaling primarily aimed to disrupt vesicular release (83), reuptake (12) or synthesis of DA (84) using knock-out animals. The DAT knock-out model suffers, however, from issues related to construct validity with respect to most DA-linked neurobehavioral disorders as homozygous loss-of-function DAT mutations in humans exhibit Juvenile Dystonia/Parkinsonism and require intensive support to live past early childhood (85,86). In the current study, we utilized mice expressing DAT Val559, an extensively characterized disease associated variant of the DAT (87), as a platform to search for a pharmacological manipulation that could normalize biochemical and behavioral traits seen with genetic DAT dysfunction. Specifically, based on considerations of the ability of KORs to regulate presynaptic mechanisms in both nigrostriatal and mesolimbic DA neurons (88,89) and recently published clinical evidence demonstrating that KOR-antagonism is well-tolerated in humans (61–65), we explored whether antagonism of KOR-mediated regulation of DAT might display therapeutic potential for the treatment of DA-linked disorders.

We report that activation of KOR enhances DAT-mediated uptake by promoting an increase in surface DAT, consistent with a previous study (54), though the prior work did not explore the post-translational modifications and motif(s)/site(s) through which KOR regulates DAT surface expression. We found that mutation of the canonical ERK1/2 site Thr53 to Ala53 prevented the KOR agonist U69,593 from inducing elevations in DA transport V_max_, DAT surface expression and phosphorylation of Thr53 DAT, providing evidence that KOR-mediated regulation of DAT is contingent on Thr53 (Figures 1 and 2).

Enhanced DAT surface expression in acute brain slices, paralleled by elevated phosphorylation at Thr53, was evident in WT VS and DS slice preparations following KOR-agonist exposure (Figure 4). Moreover, high-speed chronoamperometry in live animals revealed that while the KOR-agonist U69,593 accelerated DA clearance, the KOR-antagonist norBNI prolonged DA clearance from the extracellular space (Figure 3), establishing for the first time the dependence of *in vivo* DA clearance on endogenous KOR signaling and consistent with a link of Thr53 phosphorylation to both KOR regulated trafficking and increased DA clearance capacity. We have previously demonstrated that D2AR-mediated regulation of DAT is sex and region specific, which results in elevated DAT Val559 surface expression and Thr53 hyperphosphorylation in the DS of males (56) and VS of females (24). However, no region-specificity was observed for KOR in this regard (Figure 4) Interestingly, treatment with U69,593 significantly elevated surface DAT Val559 in the DS, without promoting a further increase in phosphorylation at Thr53 (Figure 4), which contrasts with our experiments in EM4 cells (Figure 2). This might indicate that phosphorylation at Thr53 reached a ceiling effect in the DS of DAT Val559 mice and that KOR can recruit additional and/or unknown regulatory mechanisms in native tissue preparations that are absent in heterologous expression systems. Using acute brain slices, we found that treatment with norBNI normalized the aberrant surface trafficking and hyperphosphorylation at Thr53 of DAT Val559 in the DS (56).

Mergy and colleagues have shown that expression of DAT Val559 dampens the vesicular release of DA, due to tonic stimulation of inhibitory D2ARs (18). Consequently, administration of the non-selective DAT inhibitor cocaine fails to elevate extracellular DA in the DS of male Val559 mice (20). We found that pre-treatment with norBNI fully restored the DA response to cocaine in male mice (Figure 6 and Supplementary Figure 4). This finding supports the interpretation that inhibition of KOR and subsequent normalization of surface trafficking of DAT Val559 aids in restoration of vesicular DA release in DAT Val559 mice. Additional studies are needed to clarify the contribution of somatic KOR versus presynaptic KOR in the ability of KOR antagonists to restore vesicular DA despite elevated D2AR signaling. Mice are available that lack KOR in DAergic neurons (90). Future studies with such animals should help define whether the effect of norBNI in DAT Val559 mice exclusively relies on its action at KORs expressed on DA synthesizing neurons versus other sites. Similarly, additional approaches are needed to identify the source(s) of dynorphin that allow for the ability of KOR to normalize traits in DAT Val559 mice. While a striatal source of dynorphin that could account for modulation of DAT Val559 by KORs has been documented (12), we have demonstrated that a serotonergic plasticity arises in DAT Val559 mice that drives a loss of locomotor activation by cocaine (20). In this regard, Pomrenze et al (91) have recently demonstrated that serotonergic projections to the NAc produce and release dynorphin and are worth exploring.

At present, we cannot exclude the possibility that treatment with norBNI reduces DAT Val559-mediated reverse transport *per se* as well as surface expression. DAT Val559 is hyperphosphorylated at DAT N-terminal sites that have been linked to reverse transport (42,66). Antagonism of KOR normalized the hyperphosphorylation at Thr53 and this observation could expand to other phosphorylation sites crucial for reverse transport. Further, Lycas et al (92) reported that activation of D2ARs causes DAT to accumulate in nanoclusters in which DAT appears to be biased towards the inward facing conformation. The authors found that another disease-assciated efflux-prone DAT variant (DAT-Asp421Asn) that is conformationally biased towards the inward-facing conformation preferentially localized to these clusters. Considering the tonic activation of D2ARs, DAT Val559 may reside in nanodomains that support ADE in DAergic neurons whereas antagonism of KOR could redistribute DAT Val559 into nano-environments that preclude ADE. Alternatively, antagonism of KOR could further disrupt the potentially enhanced interaction of DAT Val559 with proteins and lipids that bias DAT towards more efflux-prone states (49,93–96). Activated Gbg proteins have also been found to induce DA efflux similar to that seen with DAT Val559 and amphetamines (44,45). Whether the pathways involved with KOR antagonism intersect with this mechanism and its regulators are worthy of study. More generally, such considerations suggest that DAT mutations are likely only one mechanism by which behaviorally penetrant DA efflux states can arise, and where KOR antagonism can be of therapeutic use.

On a behavioral level, male DAT Val559 mice display a deficit in spontaneous alternation in the Y-Maze test ((24)and Figure 5), which translates into deficits in working memory (97). Administration of norBNI normalized the deficit in spontaneous alternations and direct revisits 30 min and one week post injection. The latter observation is consistent with the long-lasting effects of norBNI (77,78). Of note, a recent study demonstrated that norBNI improved working memory and attention in the perinatal nicotine exposure model, presumably via an increase in cortical DA (98). Prenatal exposure to nicotine alters development of the DA system and direct effects of this paradigm on DAT have been reported (99). Moreover, nicotinic acetylcholine receptors are well known to regulate DA vesicular release (100). This suggests that the beneficial effect of norBNI could arise from normalization of disrupted presynaptic signaling networks that dictate DA release and clearance. Future studies assessing DA dynamics in WT and DAT Val559 in mPFC, VS and DS in the context of DAT, D2AR and KOR antagonists directly administered in these regions are underway to tease apart the relative contribution of the respective terminal fields of DA neurons emanating from VTA and SNc.

In line with the persistent effect of norBNI in the Y-Maze, we also found that norBNI normalized behavior of male DAT Val559 mice in the open field one week after norBNI administration. Vehicle treated DAT Val559 mice spent significantly less time in the center than WT controls and this difference was absent when mice were treated with norBNI prior to testing (Figure 6), consistent with an anxiolytic effect. Interestingly, we (18) found no differences in horizontal locomotor activity between WT and DAT Val559 mice in the open field. The reduction in horizontal locomotor activity of male Val559 DAT mice observed in this study (Figure 6) might be attributed to repeated handling and an increased susceptibility of these mice to repeated stress. We also observed that saline treated DAT Val559 mice displayed a reduced tendency to explore the full length of the arms of the Y-Maze when they were re-exposed to the apparatus. This could indicate that the exploratory behavior is suppressed in male DAT Val559 mice upon repeated handling or when the novelty of the environment is removed. Moreover, pre-treatment with norBNI restored the locomotor response of male DAT Val559 mice to systemic cocaine in the open field (Figure 6), in agreement with re-established vesicular DA release in norBNI treated DAT Val559 mice. Of note, female DAT Val559 mice display impaired performance in the NOR task and this phenotype can be restored by systemic administration of the D2-antagonist sulpiride (24), in line with our interpretation that behavioral deficits arise from tonic activation of D2ARs. Systemic administration of norBNI improved NOR in female DAT Val559 mice (Figure 5), which suggests that antagonism of KOR rescues behavioral consequences of DAT Val559 expression regardless of sex.

In conclusion, using a construct-valid preclinical model of genetic DAT dysfunction, we found that inhibition of KOR normalizes aberrant biochemical and behavioral sequelae resulting from disrupted DAergic neurotransmission, reinforcing the potential of KOR antagonists for the treatment of disorders linked to DAT-dependent tonic DA elevation. Additionally, our findings draw attention to the interplay of D2AR and KOR signaling and their regulation of DAT and DA neurotransmission under basal conditions, as well augmentating vesicular DA release to achieve greater neuromodulatory DA “signal to noise”. Considering issues of high abuse liability with prescribing psychostimulants to treat many DA-linked psychiatric pathologies, our findings support the use of a non-stimulant type medication based on KOR antagonism for the treatment of DA-linked disorders where excess tonic vs phasic DA signaling is suspected to underlie changes in cognitive function and/or working memory.

## Acknowledgements

This work was supported by a Max Kade Fellowship of the Austrian Academy of Sciences to FPM, a NARSAD Young Investigator Grant from the BBRF to AS, P20 GM103442 to JDF, P20 GM104360 to RAV, R01 MH093320 and R21 DA046044 to LCD, R01 MH105094 to RDB and R01DA054694 to SR

