## Supplementary material for "Kappa Opioid Receptor Antagonism Restores Phosphorylation, Trafficking and Behavior induced by a Disease Associated Dopamine Transporter Variant": Suppl Information

Running Title: κ-Opioid Receptor Antagonism Normalizes DAT Val559

Felix P. Mayer, PhD[1,2,%], Adele Stewart, PhD[1,2], Durairaj Ragu Varman, PhD[3], Amy E. Moritz, PhD[4], James D. Foster, PhD[4], Anthony W. Owens[5], Lorena B. Areal, PhD[1], Raajaram Gowrishankar, PhD[1#], Michelle Velez[1], Kyria Wickham[1], Hannah Phelps[1], Rania Katamish[1], Maximilian Rabil[1], Lankupalle D. Jayanthi, PhD[3], Roxanne A. Vaughan, PhD[4], Lynette C. Daws, PhD[5,6], Randy D. Blakely, PhD[1,2]*, and Sammanda Ramamoorthy, PhD[3]*

[1] Department of Biomedical Science, Charles E. Schmidt College of Medicine, Florida Atlantic University, Jupiter, FL, USA.

[2] Stiles-Nicholson Brain Institute, Florida Atlantic University, Jupiter, FL, USA.

[3] Department of Pharmacology and Toxicology, Virginia Commonwealth University, Richmond, VA, USA

[4] Department of Biomedical Sciences, University of North Dakota School of Medicine and Health Sciences, Grand Forks, ND, USA

[5] Department of Cellular and Integrative Physiology, University of Texas Health Science Center at San Antonio, TX, USA

[6] Department of Pharmacology, University of Texas Health Science Center, San Antonio, TX, USA.

#present address: Departments of Anesthesiology and Pain Medicine and Pharmacology, University of Washington, Seattle, WA, USA and Center for the Neurobiology of Addiction, Pain and Emotion, University of Washington, Seattle, WA, USA

%present address: Department of Neuroscience, Faculty of Health and Medical Sciences, University of Copenhagen, Copenhagen, DK-2200, Denmark.

*to whom correspondence should be addressed:

Professor, Department of Pharmacology & Toxicology Virginia Commonwealth University R. Blackwell Smith Building, Room 756A P.O. Box 980613 410 North, 12th Street (at Clay St.) Richmond, VA 23298-0613, USA; Tel: 804-828-8407; Fax: 804-828-2117

Professor, FAU Stiles-Nicholson Brain Institute David J.S. Nicholson Distinguished Professor in Neuroscience, Dept Biomedical Science Charles E. Schmidt College of Medicine 5353 Parkside Drive, Jupiter, FL 33458, USA MC-22/ Room 201G, Tel: 561-799-8100

### Augmented Materials and Methods

*Materials*

Lipofectamine™ 2000, Dulbecco's Modified Eagle Medium (DMEM) and other cell culture media were purchased from Invitrogen/Life Technologies, (Grand Island, NY). [$^{3}$H]DA (dihydroxyphenylethylamine [2,5,6,7,8-3H], 63.2 Ci/mmol) and Optiphase Supermix, were purchased from PerkinElmer Inc., (Waltham, MA, USA). U69,593, U0126, protease and phosphatase cocktails were obtained from Sigma-Aldrich (St. Louis, MO). Nor-binaltorphimine dihydrochloride (norBNI) was purchased from Tocris (Bristol, United Kingdom). Reagents for SDS-polyacrylamide gel electrophoresis and Bradford protein assays were from Bio-Rad (Hercules, CA, USA), fetal bovine serum and enhanced chemiluminescence (ECL) reagents were from Thermo Fisher Scientific Inc., (Rockford, IL, USA). Sulfosuccinimidyl-2-[biotinamido] ethyi-1,3-dithiopropionate (EZ link NHS-Sulfo-SS-biotin), Protein A magnetic beads (Dynabeads) and NeutrAvidin Agarose were purchased from Thermo Scientific (Waltham, MA, USA). Anti-Calnexin (Cat# ADI-SPA-860-D, RRID: AB-10616095) was obtained from Enzo Life Sciences, Inc (Farmingdale, NY, USA). QuikChange II XL site-directed mutagenesis kit was obtained from Agilent Technologies (Santa Clara, CA, USA). Peroxidase-affinipure goat anti-rabbit IgG antibody (HRP conjugated secondary antibody (Cat# 111-035-003, RRID: AB-2313567) was acquired from Jackson Immuno Research Laboratories (West Grove, PA). DAT antibody Cat# 431-DATC, RRID: AB-2492076 – was used for experiments using EM4 cells. DAT antibody MAB369 (Millipore Sigma, RRID:AB_2190413) -was used for experiments using acute slices). DAT antibody MABN669 (Millipore Sigma, RRID:AB_2717269) was used for experiments in rat striatal synaptosomes. DAT Thr53 Antibody p435-53 (PhosphoSolutions, RRID: AB-2492078) was used to detect DAT phosphorylated at Thr53 in all experiments.

*Cell culture and transfection of WT and mutant cDNAs*

EM4 cells were maintained in DMEM media with 10% fetal bovine serum, 1% penicillin-streptomycin and 1% glutamine as described previously (1). For DA transport assays, EM4 cells were seeded onto 24 well plates (100,000 cells/well). For surface biotinylation experiments, 150,000 cells/well were seeded onto 12 well plates. After 24 hrs of seeding, cells were co-transfected with cDNA constructs encoding myc-KOR plus rDAT-WT, or myc-KOR plus rDAT Ala53, or myc-KOR plus pcDNA3 using Lipofectamine 2000 according to the manufacturer's instruction. QuikChange II XL site-directed mutagenesis kit was used to alter rat DAT cDNA sequence encoding alanine at amino acid 53 (Ala53) to encode threonine (Thr53). Sequence validation of the substitution and absence of nonspecific mutations were confirmed by sequencing the entire cDNA on both strands.

The following DNA amounts were used for transfections: myc-KOR:0.25 µg/well (24 well plate) 0.5 µg/well (12 well plate); rDAT, or rDAT Ala53: 0.25 µg/well (24 well plate) 0.5 µg/well (12 well plate). Matched amounts of pcDNA3 were transfected for control experiments. The plates were further incubated in a humidified atmosphere at 37˚C and 5% $CO_2$ for 24 hrs prior to assay.

*$[^3H]$DA uptake assay*

EM4 cells co-transfected with KOR plus rDAT, or rDAT Ala53, or pcDNA3 in 24 well plates were first washed with serum free DMEM (1 mL/well) media and incubated at 37°C for 2 hrs. Cells were then washed with 1 mL of pre-warmed (37˚C) modified Krebs-Ringer-HEPES (KRH) buffer, pH 7.4 supplemented with L-ascorbic acid and pargyline (10 mM HEPES, 120 mM NaCl, 4.7 mM KCl, 1.2 mM $KH_2PO_4$, 1.2 mM $MgSO_4$, 10 mM D-glucose, 0.1 mM L-ascorbic acid, 0.1 mM pargyline, pH 7.4). Modulators were added for the times noted in figures and/or legends, followed by the addition of 30 nM $[^3H]$DA to initiate DA uptake. $[^3H]$DA uptake was terminated after 5 min of incubation at 37˚C by removing the uptake mixture and rapidly washing with 1.0 mL KRH buffer three times. Cells were lysed by adding 0.2 mL Optiphase Supermix cocktail and accumulated $[^3H]$DA-radioactivity was measured using a microplate scintillation counter (MicroBeta$^2$ LumiJET, PerkinElmer, Waltham, MA, USA). DAT-specific $[^3H]$DA transport was calculated as the difference in counts obtained in the presence or absence of the DAT-specific antagonist GBR12909 (100 nM). The effectiveness of GBR12909 in blocking DAT activity was verified using cells transfected with pcDNA3 vector alone. For DAT kinetic analyses, unlabeled DA was mixed with 30 nM $[^3H]$DA, with the final concentration of DA varying from 0.25 µM to 20 µM. Uptake assays were performed in triplicate.

*EM4 cell surface biotinylation*

Cell surface biotinylation was performed as described previously (1). EM4 cells co-transfected with KOR plus rDAT, or rDAT Ala53, or pcDNA3 in 12 well plates were washed and preincubated with serum free DMEM for 2 hrs followed by treatment with vehicle or U69,593 (10 µM) for 30 min. Biotinylation of cell surface proteins was initiated by incubating cells with EZ link NHS-Sulfo-SS-biotin (1 mg/mL) in cold PBS/$Ca^{2+}$-$Mg^{2+}$ buffer (138 mM NaCl, 2.7 mM KCl, 1.5 mM $KH_2PO_4$, 9.6 mM $Na_2HPO_4$, 1 mM $MgCl_2$, 0.1 mM $CaCl_2$, pH 7.3) for 30 min on ice followed by removal of excess biotinylating reagents by washing the cells twice with ice-cold PBS/$Ca^{2+}$-$Mg^{2+}$ buffer containing 100 mM glycine and incubated further with glycine (100 mM) for 20 min on ice to quench excess NHS-Sulfo-SS-biotin. The cells were solubilized with 0.7 mL/well of RIPA buffer (10 mM Tris-HCl, pH 7.5, 150 mM NaCl, 1 mM EDTA, 1% Triton X-100, 0.1% SDS, 1% sodium deoxycholate) containing a protease and phosphatase inhibitor cocktail. Surface biotinylated proteins were isolated from equal amounts of solubilized proteins using

NeutrAvidin Agarose resins by overnight incubation at 4˚C followed by washing with RIPA. Bound proteins were eluted in 50 µL Laemmli sample buffer (62.5 mM Tris-HCl pH 6.8, 20 % glycerol, 2 % SDS and 5 % ß-mercaptoethanol) by incubating for 30 min at room temp. All eluates and aliquots (40 µl) of total extracts and unbound fractions were used for SDS-PAGE and immunoblot analysis as described below under immunoblot analysis. Proteins were separated using 7.5% SDS-polyacrylamide gels followed by transfer to polyvinylidene difluoride membranes (PVDF, ThermoFisher Scientific). To analyze total DAT and calnexin proteins, membranes were pre-blocked with PBS containing 0.1% Tween 20 and 1% nonfat milk, whereas Tris buffer containing 1% BSA and 0.1% Tween 20 was used for the detection of Thr53-pDAT. Membranes were incubated with primary antibodies to DAT or Thr53-pDAT (1: 1000) overnight at 4˚C. After washing, secondary antibody peroxidase-affinipure goat anti-rabbit IgG antibody (1: 10,000) was incubated for 1 hr at room temp followed by detection of immunoreactive proteins using ECL or ECL plus reagent (ThermoFisher Scientific). Using NIH Image J 1.52a, bands were quantified from digitized blots within the linear range of the film exposure. Specific nature of DAT bands was defined by the absence of corresponding immunoreactive signal in pcDNA3 transfected cells analyzed in parallel. Blots were also stripped and reprobed with calnexin antibody(1:10000 dilutions) to normalize total DAT and p-Thr53 DAT.

*Animals*

All procedures involving animals were approved by the Institutional Animal Care and Use Committees of UT San Antonio Health Sciences Center, Virginia Commonwealth University, or Florida Atlantic University depending on the site of assays, in accordance with the National Institutes of Health Guide for the Care and Use of Laboratory Animals. Male Sprague-Dawley rats (175-350 g body weight) were utilized for *in vivo* chronoamperometry and assays of DAT Thr53 phosphorylation in synaptosomes and tissue. For slice experiments, age-matched 4- to 6-week-old male homozygous mice for either WT or DAT Val559 (genetic background: 75% 129/6 and 25% C57; (2)) were bred from homozygous dams and sires that were derived from heterozygous breeders. Male mice were used due to the male bias observed in ADHD diagnosis (3). All biochemical experiments were conducted during the light phase. For behavioral experiments, 6–8-week-old WT and homozygous DAT Val559 littermates derived from heterozygous breeders were utilized. Male mice were used for all behavioral experiments except the novel object recognition test due to the demonstrated sex bias of behavioral phenotypes observed in DAT Val559 mice (4). Behavioral testing was performed during the dark phase of the light cycle under red light. Rodents were maintained in a temperature and humidity-controlled room on a 12:12 h light/dark cycle. Food and water were supplied *ad libitum*. All efforts and care were taken to minimize animal suffering and to reduce the number of animals used. As alternatives to brain tissues, cell culture models were utilized.

*Treatment of striatal synaptosomes with U69,593 ± U0126*

Male Sprague-Dawley rats (175–300 g) were decapitated, and striata were rapidly dissected, weighed, and placed in ice-cold sucrose phosphate (SP) buffer (0.32 M sucrose and 10 mm sodium phosphate, pH 7.4). Tissues were homogenized in ice-cold SP buffer with 15 strokes in a glass/Teflon homogenizer and centrifuged at 3000 × *g* for 3 min at 4 °C. Supernatant fractions were re-centrifuged at 17,000 × *g* for 12 min, and the resulting P2 pellet enriched for synaptosomes was resuspended to 20 mg/mL original wet weight in ice-cold SP buffer. Synaptosomes were treated with vehicle or 50 μM U0126 for 15 min at 30˚C, followed by treatment with vehicle or 10 μM U69,593 for an additional 15 min at 30˚C. After treatment, samples were subjected to SDS-PAGE followed by immunoblotting for total DAT and DAT phosphorylated at Thr53 (p-Thr53 DAT levels).

*p-Thr53 DAT levels from rats injected with vehicle or U69,593*

Male rats were injected s.c. with vehicle or 0.32 mg/kg U69,593 and sacrificed at the indicated times noted in Figure 3. DS or VS were dissected, weighed and kept frozen until analyzed. The pre-weighed samples were homogenized with a Polytron PT1200 homogenizer (Kinematica, Basel, Switzerland) for 8 s in ice-cold SP buffer, and centrifuged at 3000 *× g* for 3 min at 4 °C. Supernatant fractions were re-centrifuged at 17,000 *× g* for 12 min to obtain a membrane pellet. The pellet was resuspended to 20 mg/mL original wet weight in ice-cold SP buffer. Samples were subjected in duplicate to SDS-PAGE and western blot for total and p-Thr53 DAT, and p-Thr53 DAT staining quantified by normalization to the time-matched controls.

*Analysis of in vivo DA clearance in NAc*

High-Speed *in vivo* chronoamperometry, conducted as previously described (5), was used to determine the effect of KOR modulators on DA clearance in NAc of anesthetized rats. Briefly, carbon fiber electrodes were attached to multi-barrel glass micropipettes. The tips of carbon fibers (150 μm in length) were coated with 5% Nafion to provide a 1000-fold selectivity of DA over its metabolite dihydroxyphenylacetic acid (DOPAC) and displayed linear amperometric responses to 0.5 – 10 μM DA during *in vitro* calibrations. Rats were anesthetized with chloralose (85 mg/kg, i.p.) and urethane (850 mg/kg, i.p). Electrodes were lowered into the NAc (Anterior/Posterior, +1.7 mm relative to bregma; medial/lateral, ±0.9 mm relative to bregma; V, -6.0 mm from dura; (6)using a stereotaxic frame. Throughout the experiments, body temperature was maintained at 37 ±1°C by a water-circulated heating pad. Individual micropipette barrels were filled with DA (200 μM), U69,593 (885.7 μM, barrel concentration), norBNI (400 μM, barrel concentration) and vehicle and ejected (100–150 nL) using a Picospritzer II (General Valve Corporation, Fairfield, NJ), to deliver 89 to 134 pmol U69,593, or 40 to

60 pmol norBNI. Note that there is a 10-200 fold dilution in the concentration of drug reaching the area surrounding the recording electrode (5), thus concentrations of U69,593 and norBNI at the recording electrode are estimated to range from 5 to 89 µM, and 2 to 40 µM, respectively. Oxidation potentials consisted of 100 ms pulses of +0.55 V alternated with 100 ms intervals during which the resting potential was maintained at 0 V. Oxidation and reduction currents were digitally integrated during the last 80 ms of each 100 ms voltage pulse. DA was pressure-ejected until a stable baseline signal was established (typically after 3-4 ejections). The effect of U69,593, norBNI, and vehicle on DA signals was quantified 2 min after a stable baseline DA signal was established. Changes in T50 (time required for the signal to decline by 50% of its peak amplitude) and T80 (time required for the signal to decline by 80% of its peak amplitude) were used to determine the influence of U69,595 (KOR activation) and norBNI (KOR inhibition) or vehicle on DA clearance.

*Surface biotinylation and immunoprecipitation of p-Thr53 DAT using acute mouse brain slices*

All procedures were performed as described previously (Gowrishankar et al., 2018).

*Acute slice preparation:* Male 4-6 week-old mice were sacrificed by rapid decapitation, brains were removed and transferred into ice-cold oxygenated sucrose artificial cerebrospinal fluid (sucrose-aCSF; in mM: sucrose 250, KCl 2.5, $NaH_2PO_4$ 1.2, $NaHCO_3$ 26, D-glucose 11, MgCl2.6$H_2O$ 1.2, $CaCl_2.2H_2O$ 2.4, pH 7.4, 300 –310 mOsm), Subsequently, 300 µm thick coronal slices (Anterior/Posterior: 1.5-1.0 mm for the ventral striatum; 1.0-0.2 mm for the dorsal striatum) were collected using a Leica VT100 S vibratome in ice-cold oxygenated sucrose-aCSF. Slices were then allowed to recover at 30-32 °C for 45-60 min in standard aCSF (in mM: NaCl 124, KCl 2.5, $NaH_2PO_4$ 1.2, $NaHCO_3$ 26, D-glucose 11, $MgCl_2.6H_2O$ 1.2, $CaCl_2.2H_2O$ 2.4, pH 7.4, 300 –310 mOsm) with constant oxygenation. Afterwards, slices were transferred to a water bath kept at 37 °C and washed three times with standard aCSF (5 min each) before drugs or vehicle were added (U69,593: 10 µM for 7 min. norBNI: 1 µM for 20 min) with constant oxygenation. After drug treatments, the slices were subjected to three rapid washes and two 5-min washes with ice cold oxygenated standard aCSF. For immunoprecipitation studies, VS and DS were dissected at this point and tissue was flash frozen in liquid nitrogen and stored at -80 °C until processed. For biotinylation experiments, the slices were exposed to 1 mg/mL EZ link NHS-Sulfo-SS-biotin for 30 min on ice with constant oxygenation. Subsequently, the reaction was quenched by exposing the slices to 0.1 M glycine (in standard-aCSF; two 10 min incubations). Finally, slices were washed with ice-cold standard-aCSF (three rapid and two 5-min washes) and ventral and dorsal striatum were dissected and transferred into RIPA buffer (in mM: Tris-HCl 25 pH 7.6, NaCl 150, EDTA 5, 1% Triton X-100, 1% sodium deoxycholate, 0.1% SDS) supplemented with protease inhibitor cocktail (P8340, Millipore Sigma; used at a 1:100 dilution). Tissue was solubilized by passing the slices through a 29 gauge needle and nutation at 4 °C for 60 min. Subsequently, samples were centrifuged at 4 °C for 15 min at 21,000 *x g*, pellets were discarded, and supernatants transferred into fresh tubes. Subsequently, protein concentrations were determined using the BCA protein assay (ThermoFisher) and assessed with a FLUOstar Omega microplate reader (BMG Labtech). Bovine serum albumin was used as standard. Lysates were exposed to Pierce™ Streptavidin Agarose beads (ThermoFisher; 20353) (ratio: 20 µg of protein to 50 µL bead slurry) (minimum 2 hrs, maximum 12 hrs) under constant nutation. Beads were washed 3 times with RIPA buffer and protein was eluted and denatured by adding 2X sample buffer (0.5 mM Tris-HCl pH 6.8, 25% glycerol, 1.0% bromophenol blue, 10% SDS, 5 % beta-mercaptoethanol) at room temp for 30 min on a shaker. 5 µg of total protein was loaded on the same gel to normalize the surface DAT to total DAT present in each sample.

*Immunoprecipitation:* Slices were transferred into ice-cold lysis buffer (in mM: NaCl 138, KCl 2.7, TRIS 50, 1% Triton X-100, pH 8) supplemented with protease (P8340, Millipore Sigma) and phosphatase (P0044, Millipore Sigma) inhibitors (both used at a 1:100 dilution). Tissue was solubilized by carefully passing the slices through a 29 Gauge needle and subsequent nutation at 4 °C for 60 min. Samples were centrifuged at 4 °C for 15 min at 21000 *x g*, pellets were discarded and supernatants transferred into fresh tubes. Rabbit Thr53 DAT antibody (RRID:AB_2492078) was crosslinked to protein A magnetic beads (Dynabeads, ThermoFisher; ratio 1 µL Thr53 antibody:10 µL Dynabeads as described (7). 20 µL of DAT Thr53 antibody-conjugated beads were added to 200 µg of protein and nutated (minimum 4 hrs, maximum 10 hrs) at 4 °C. Subsequently, beads were washed with lysis buffer (4 times) and incubated with 50 uL of 2X sample buffer (0.5 mM Tris-HCl pH 6.8, 25% glycerol, 1.0% bromophenol blue, 10% SDS, 5 % beta-mercaptoethanol) for 45-60 min at room temp on a shaker to elute/denature protein before SDS-PAGE and immunoblotting for DAT. 5 µg of total protein was loaded on the same gel to normalize the immunopurified p-Thr53 DAT to total DAT present in each sample.

*Surface biotinylation following in vivo drug treatment:* 4-6 week old male mice were injected with norBNI (10 mg/kg, i.p. or saline. Mice were sacrificed 30min post drug administration by rapid decapitation, brains were removed and transferred into ice-cold oxygenated sucrose-aCSF, Subsequently, 300 µm thick coronal slices (Anterior/Posterior: 1.5-1.0 mm for the ventral striatum; 1.0-0.2 mm for the dorsal striatum) were collected using a Leica VT100 S vibratome in ice-cold oxygenated sucrose-aCSF. Slices were then washed three times with ice-cold standard aCSF and exposed to 1 mg/mL EZ link NHS-Sulfo-SS-biotin for 30 min on ice with constant oxygenation. Subsequently, the reaction was quenched by exposing the slices to 0.1 M glycine (in standard-aCSF;

two 10 min incubations). Finally, slices were washed with ice-cold standard-aCSF (three rapid and two 5-min washes) and ventral and dorsal striatum were dissected and transferred into RIPA buffer (in mM: Tris-HCl 25 pH 7.6, NaCl 150, EDTA 5, 1% Triton X-100, 1% sodium deoxycholate, 0.1% SDS) supplemented with protease inhibitor cocktail (P8340, Millipore Sigma; used at a 1:100 dilution). Tissue was solubilized by passing the slices through a 29 gauge needle and nutation at 4 °C for 60 min. Subsequently, samples were centrifuged at 4 °C for 15 min at 21,000 *x g*, pellets were discarded, and supernatants transferred into fresh tubes. Subsequently, protein concentrations were determined using the BCA protein assay (ThermoFisher) and assessed with a FLUOstar Omega microplate reader (BMG Labtech). Bovine serum albumin was used as standard. Lysates were exposed to Pierce™ Streptavidin Agarose beads (ThermoFisher; 20353) (ratio: 20 µg of protein to 50 µL bead slurry) (minimum 2 hrs, maximum 12 hrs) under constant nutation. Beads were washed 3 times with RIPA buffer and protein was eluted and denatured by adding 2X sample buffer (0.5 mM Tris-HCl pH 6.8, 25% glycerol, 1.0% bromophenol blue, 10% SDS, 5 % beta-mercaptoethanol) at room temp for 30 min on a shaker. 5 µg of total protein was loaded on the same gel to normalize the surface DAT to total DAT present in each sample.

*Behavioral testing*

Y Maze and Open Field Testing – On day 1, 6–8-week-old WT and homozygous DAT Val559 male mice received a single injection of saline or norBNI (10 mg/kg, i.p.) 30 mins prior to behavioral testing and were returned to their home cage until experiment initiation. Animals were subsequently placed in a Y-shaped maze combining 3 white, opaque plastic arms (14 × 4.5 × 40 cm) each at a 120°C angle from the others. Each subject was introduced to the center of the maze and allowed to freely explore all arms for 10 min. Locomotor activity and arm entries were monitored using Ethovision XT video tracking software (version 15; Noldus, Leesburg, VA) and the percent alternation calculated as the number of alternations (incidences where animals visited each arm in turn without repeating an arm) divided by the total number of arm visit triads X 100. 1 week later, animals were again placed in the maze to assess whether drug and genotype effects were maintained in accordance with the long lasting effects of norBNI (8,9). 2 days after the second Y Maze test, the same animals were placed in an open field apparatus (27 cm X 27 cm x 20.5 cm) contained within light- and air-controlled sound-attenuating boxes (Med Associates, Fairfax, VT). As described previously (4), locomotion (horizontal/vertical) as well as stereotypic behaviors and center occupancy were detected by interruption of a grid of infrared beams (16 photocells in each horizontal axis and elevated 4 cm above the floor). Data were collected and analyzed by Med Associates Activity Monitor software.

Cocaine-induced locomotor activation - To assess the effect of norBNI on cocaine-induced locomotor activity, 6–8-week-old WT and homozygous DAT Val559 DAT male mice were injected with norBNI (10 mg/kg, i.p.) one week prior to the open field test and then received either saline or cocaine (Sigma, 10 mg/kg, i.p.) immediately prior to placement in Med Associates open field apparatuses. Locomotor activity was collected for 60 minutes following drug injection utilizing Med Associates Activity Monitor software as above.

Novel Object Recognition (NOR) test - The novel object recognition (NOR) test was performed as described in Stewart et al., 2022 (4). Female WT and DAT Val559 mice (6-8 weeks of age) were utilized for this test as the behavior of DAT Val559 males is unaltered relative to WT littermates (4). Mice were habituated to the test apparatus for 10 min once per day for 2 days. On day 3, mice received a single injection of saline or norBNI (10 mg/kg, i.p.) 30 mins prior to placement in the test apparatus with two identical objects. Given that norBNI impacts are long lasting, mice were not re-dosed on day 4 when they were allowed to freely explore a familiar object or novel object. Activity was monitored with Ethovision XT software (version 15) and data are presented as % time exploring each object or discrimination index [(time with novel object – time with familiar object)/total exploration time].

*In vivo microdialysis*

Surgeries to insert microdialysis guide cannulae were performed as described earlier (2,10). 6–8-week-old male mice were anesthetized using isoflurane (5 % induction/ 2 % maintenance) and their heads were fixed in a stereotaxic frame. Ophthalmic ointment was applied to prevent drying of the eyes. Analgesics: bupivacaine and lidocaine were administered locally (100 µL of sterile saline containing 0.05% bupivacaine and 0.2% lidocaine, subcutaneous injection (s.c.)) and ketoprofen (10 mg/kg, s.c.) was administered systemically. Following a midline incision, lambda and bregma were levelled and a craniotomy was performed using a dental drill. Microdialysis was performed as described in Mergy et al., 2014 (2), with minor modifications: a 5 mm guide cannula (S-5000; Synaptechnology Inc., Marquette, MI) was lowered into the DS using the following target coordinates (referring to the tip of the guide cannula): Anterior/Posterior 0.86 mm, Medial/Lateral 1.6 mm, Dorsal/Ventral - 2.3 mm (all relative to bregma). The guide cannula was affixed to the skull with dental cement, supported by three 1.6 mm screws (Plastics One Inc., 00-96X1/16 39052; Fisher Scientific). Subsequently, mice were allowed to recover for 6 days. 12 hr prior to the experiment a microdialysis probe with an active membrane length of 3 mm (S-5030; Synaptechnology Inc., Marquette, MI) was inserted into the guide cannula and mice were placed into 15 inch high, plexiglass recording chambers (MTANK W/F (Instech, Plymouth Meeting, PA, USA)) with bedding and food and water provided *ad libitum*. The probe was superfused with artificial cerebrospinal fluid (in mM: NaCl 149, 2, KCl 2.8, $CaCl_2$ 1.2, $MgCl_2$ 1.2, and D-glucose 5.4, pH 7.2) overnight (12h) at a flow rate of 1 µL per min before the collection of dialysates was initiated. 20 min fractions were collected on ice and stored at -80 °C until they were analyzed with HPLC-EC (Neurobehavior Core, FAU Stiles-Nicholson Brain Institute). After collection of the three (0-60 min) baseline fractions, mice received an i.p. injection of norBNI (10 mg/kg). After four additional fractions, cocaine hydrochloride (10 mg/kg; i.p.) was injected. Changes in extracellular DA were expressed as fold increase of basal DA (i.e. the average DA of the first three basal dialysates).

*Statistical analyses*

Prism 7 (GraphPad, San Diego, CA) was used for data analysis and graph preparation. Values are presented as mean ± standard deviation (SD). One-way or two-way ANOVA were used followed by post hoc testing for multiple comparisons. The type and results of post hoc tests are noted in the Figure legends. Two-tailed, unpaired Student's t-tests were performed for comparisons between two groups. P values ≤ 0.05 were considered statistically significant.

*Data Availability*

All data will be provided by the corresponding authors upon request.

### Supplementary Figures and Legends

#### Supplementary Figure 1

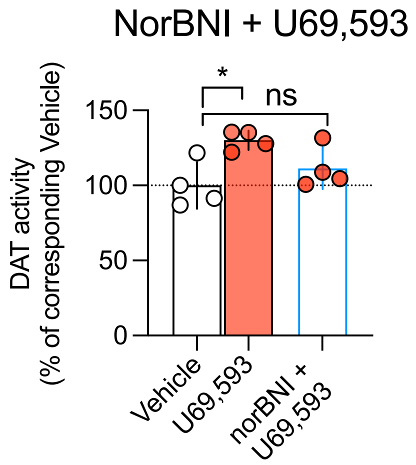


**Supplementary Figure 1: The KOR antagonist norBNI eliminates stimulation of rDAT activity by U69,593**

EM4 cells stably coexpressing rDAT and rKOR were exposed to either vehicle, U69,593 (10 µM, 15 min) or u69,593 (10 µM) plus norBNI (1 µM) (15 min) prior to the addition of [$^3$H]DA (30 nM) with specific uptake (i.e. uptake sensitive to the DAT antagonist GBR12909) assessed over 5 min. Exposure to U69,593 alone significantly increased specific uptake of [$^3$H]DA whereas treatment of U69,593 in the presence of norBNI yielded no significant stimulation vs vehicle treatments. Data are shown as mean and SD of n=4 experiments. Data were analyzed using one-way ANOVA, Bonferroni's multiple comparisons test; *= P≤0.05, ns = not significant.

### Supplementary Figure 2

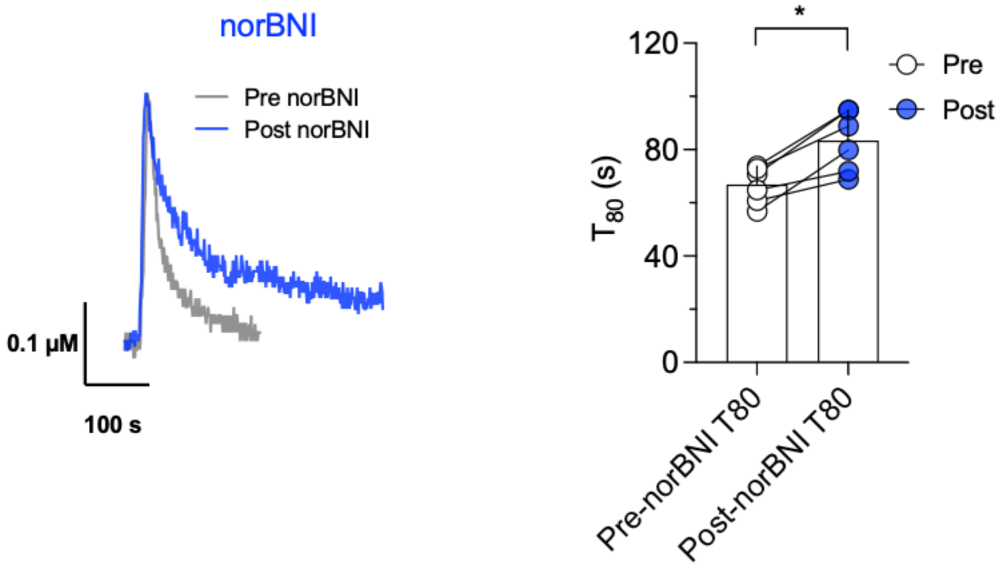


**Supplementary Figure 2: norBNI increases clearance time of DA in NAc**
Left: representative oxidation currents converted to micromolar values that were recorded upon pressure ejection of DA before (gray trace) or after injection of norBNI (blue trace) into NAc. Right: norBNI significantly increased the amount of time that was required to clear 80% of the peak signal. Individual values from each animal are reflected by the corresponding symbols. *= $P<0.05$ (two-tailed t-test, n=6 per group). Bars and error bars represent the mean and SD.

### Supplementary Figure 3

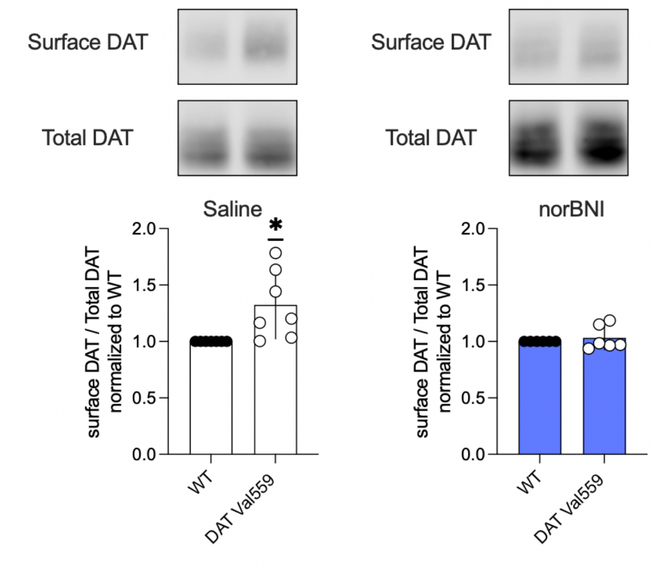


**Supplementary Figure 3: Systemic treatment with norBNI normalizes DAT Val599 surface expression**

Male WT and Val559 mice were injected with vehicle (saline, n=7 for both genotypes) or norBNI (10 mg/kg, i.p., n=6 for both genotypes) 30 min before acute coronal slices containing the DS were collected and subjected to immediate surface biotinylation. Biotinylated proteins were extracted and surface DAT was compared to total DAT levels. Data are shown as mean and SD. *=P<0.05, one-sample t-test (two-tailed).

### Supplementary Figure 4

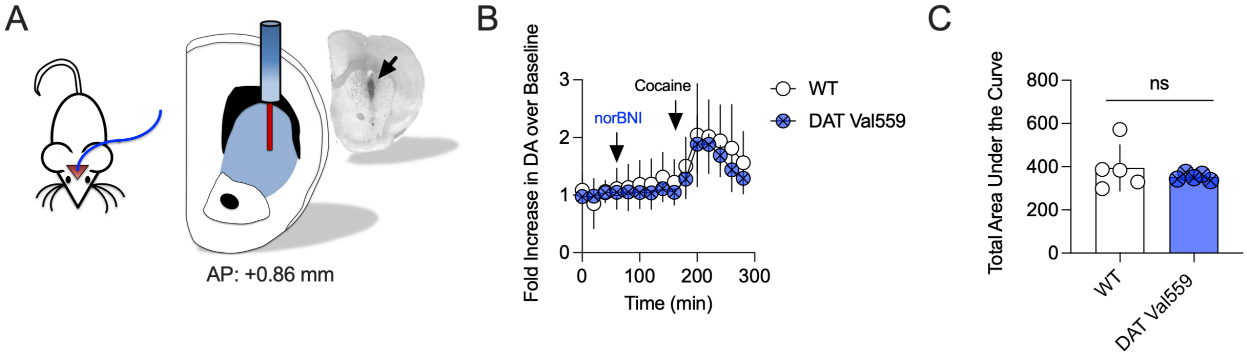


**Supplementary Figure 4: Pretreatment with norBNI restores cocaine-induced increase in extracellular DA the DS of freely moving DAT Val559 mice**

**A)** Schematic displaying the placement of the microdialysis probe in the DS. The insert shows a coronal slice with the black arrow indicating the position of the active dialysis probe.

**B)** Male WT and DAT Val559 mice were injected with norBNI (10 mg/kg, i.p.) and cocaine (10 mg/kg, i.p.) and extracellular DA was measured using *in vivo* microdialysis in the DS.
20 min samples were collected and DA is expressed as fold increase of basal DA (i.e. the average from t=0 to 60 min). Arrows indicate times for systemic administration of norBNI and cocaine. N=5 animals per genotype, data are shown as mean and SD.

**C)** Shown is the total area under the curve for the traces shown in **(B)**. Data are shown as the mean and error bars indicate the SD. Ns= not significant, unpaired Student's t-test with Welch's correction.

### Supplementary Figure 5

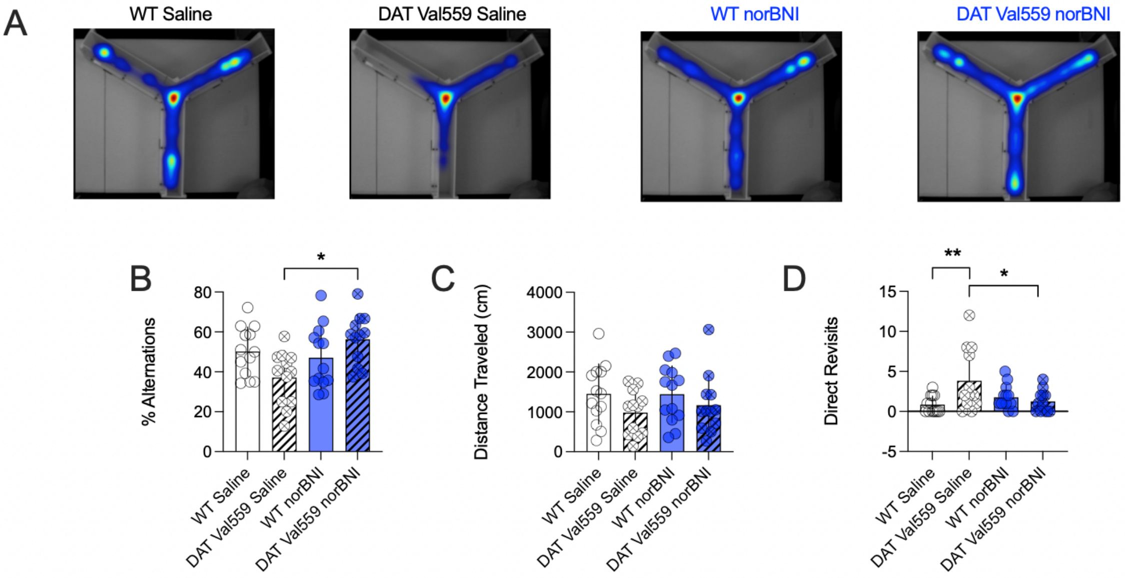


**Supplementary Figure 5: Pretreatment with norBNI one week prior to testing normalizes the repetitive explorative behavior of male Val599 mice in the Y-Maze test.**
The same cohort of mice shown in Figure 5 was retested in the Y-Maze one week after the administration of norBNI with representative heat maps shown in A).
B) DAT Val559 mice treated with vehicle displayed significantly fewer alternations when compared to DAT Val559 mice injected with norBNI.
C) No differences between genotypes or drug treatment groups on total distance travelled were detected one week after norBNI administration.
D) Treatment with norBNI restored direct revisits of DAT Val559 mice to levels that were comparable to the WT controls.
All bars show the mean and SD. n=12-13 individual animals per group, data in B, C and D were analyzed with two-way ANOVA, Šídák's multiple comparisons test. Symbols reflect individual animals. * = $P \leq 0.05$, ** = $P \leq 0.01$.
